# Trypanosoma brucei Mitochondrial DNA Polymerase POLIB Contains a Novel Thumb Insertion That Confers Dominant Exonuclease Activity

**DOI:** 10.1101/2022.08.12.503786

**Authors:** Stephanie B. Delzell, Scott W. Nelson, Matthew P. Frost, Michele M. Klingbeil

## Abstract

*Trypanosoma brucei* and related parasites contain an unusual catenated mitochondrial genome known as kinetoplast DNA (kDNA) composed of maxicircles and minicircles. The kDNA structure and replication mechanism are divergent and essential for parasite survival. POLIB is one of three Family A DNA polymerases independently essential to maintain the kDNA network. However, the division of labor among the paralogs, particularly which might fulfill the role of a replicative, proofreading enzyme remains enigmatic. *De novo* modeling of POLIB revealed a structure divergent from all other Family A polymerases in which the thumb subdomain contains a 369 amino acid insertion with homology to DEDDh DnaQ family 3’-5’ exonucleases. Here we demonstrate recombinant POLIB 3’-5’ exonuclease prefers DNA vs. RNA substrates and degrades single- and double-stranded DNA in a non-processive manner. The exonuclease activity prevails over polymerase activity on DNA substrates at pH 8.0, while DNA primer extension is favored at pH 6.0. Mutations that ablate POLIB polymerase activity slow the exonuclease rate suggesting crosstalk between the domains. We show that POLIB is able to extend an RNA primer more efficiently than a DNA primer in the presence of dNTPs but does not incorporate rNTPs efficiently using either primer. Immunoprecipitation of Pol I-like paralogs from *T. brucei* corroborate the pH selectivity and RNA primer preferences of POLIB and revealed that the other paralogs efficiently extend a DNA primer. The unique POLIB thumb insertion influences the balance between polymerase and exonuclease activity and provides another example of exquisite diversity among DNA polymerases for specialized function.

## INTRODUCTION

Family A polymerases (pols) have a wide variety of functions in DNA replication and repair across all domains of life. Some of these pols, including *Homo sapiens* θ and *Escherichia coli* Pol I are critical to DNA repair pathways. In eukaryotes, organellar DNA replication is also performed by Family A DNA pols. Examples include *H. sapiens* Pol g and *Saccharomyces cerevisiae* Mip1 which are replicative DNA pols for mitochondrial DNA. In *Arabidopsis thaliana,* two Family A pols (At PolIA and AtPolIB) replicate both the chloroplast and mitochondrial DNA. Replication in these organelles likely uses a scheme where one of these proteins, PolIA, is the replicative enzyme while PolIB plays a supporting repair role.^1^ Even in divergent systems of organellar DNA replication, such as within the quadruple-membrane bound organelle, the apicoplast, found in the malaria-causing protist *Plasmodium falciparum,* a Family A polymerase is the replicative enzyme (PREX).^2^

Another example of a highly divergent organellar DNA replication system is found in the group of protists named trypanosomatids. This group includes several species of parasites that are notorious human pathogens, including *Leishmania donovani* which causes Leishmaniasis, *Trypanosoma cruzi* which causes Chagas disease, and *Trypanosoma brucei* which causes African sleeping sickness in humans and Nagana in cattle. Trypanosomatids are characterized by their mitochondrial DNA structure, kinetoplast DNA (kDNA). The kinetoplast is composed of catenated circular kDNA molecules in two categories: minicircles and maxicircles. In the well-studied trypanosomatid organism, *Trypanosoma brucei,* minicircles are 1 kb each and maxicircles are 23 kb.^3^ These circular DNA molecules are condensed into a disc-shaped network found in close proximity to the flagellar basal body of these organisms.

In *T. brucei,* six DNA polymerases localize exclusively to the mitochondrion. These polymerases include two paralogs belonging to Family X, Pol β and Pol β-PAK and four Family A Pol I-like paralogs, POLIA, POLIB, POLIC, and POLID.^4,5^ This multiplicity of DNA pols is likely the result of a gene duplication event since the four Pol I-like paralogs are conserved in all kinetoplastid organisms sequenced to date.^6,7^ RNAi studies have shown that three of the Pol I-like proteins, POLIB, POLIC, and POLID, are essential for successful replication of the complex kDNA nucleoid found in the mitochondrion.^5,8,9^

RNAi of POLIB or POLID resulted in loss of fitness, disrupted the balance of minicircle replication intermediates, and caused progressive loss of the kDNA network; hallmarks of a kDNA replication defect. Yet, a role in maxicircle replication was never addressed. Structure function studies of POLIC using RNAi complementation revealed that POLIC is a dual-functioning DNA Pol with essential roles in nucleotidyl incorporation and a non-catalytic role in kDNA distribution.^10^ However, the precise division of labor among the paralogs, particularly which protein(s) is a processive, proofreading enzyme at a replication fork remains enigmatic.

Family A DNA polymerases have a conserved structure resembling a right hand with subdomains labeled, fingers, palm, and thumb. This structure of Family A DNA polymerases are often compared to the founding member of the family, *E. coli* Pol I. Variations found in other Family A DNA pols include insertions that modulate their functions. An insertion in T7 DNA pol allows for binding to *E. coli* thioredoxin and enhanced processivity of the enzyme.^11^ In plant organellar polymerases (POPs), small insertions allow for the enzymes to perform translesion synthesis.^12^ Human mitochondrial DNA polymerase (Pol γ) has a 310 amino acid spacer region in its thumb subdomain that improves the intrinsic processivity of the enzyme and allows for it to bind to the non-catalytic subunit.^13^

Through *de novo* protein structure prediction, we demonstrated an insertion in the thumb subdomain of POLIB, larger than previously described insertions at 369 amino acids in length. This insertion houses the catalytic residues typical of an exonuclease (exo) domain. Using recombinant purified protein, we confirmed this domain is indeed an active 3’-5’ exonuclease that is capable of degrading both DNA and RNA oligos, with a preference for single stranded substrates. We also demonstrate nucleotidyl incorporation activity of POLIB for the first time and demonstrate a lower pH tolerance for this activity than for POLIB exo activity. POLIB also incorporates nucleotides more rapidly from an RNA primer than a DNA primer, suggesting an RNA primed template may be its more natural substrate. Our data suggest that POLIB is likely not contributing to kDNA replication as the replicative polymerase but is contributing to nucleic acid metabolism in the mitochondrion through exonuclease activity and short extension from an RNA primer.

## MATERIALS AND METHODS

### Materials

All chemicals were molecular biology grade or better. Ultrapure NTPs were from New England Biolabs and dNTPs were from Invitrogen. Primers containing ribonucleotides, fluorescein (6-FAM), or a 2-aminopurine (2AP) were synthesized by Integrated DNA Technologies. 5**’** hexachlorofluorescein (HEX)-labeled RNA primer was gel-purified prior to use. All other primers were synthesized by Invitrogen. A full listing of primers used in this study can be found in Table 1, Table S1 and Table S2.

**Table 1.**
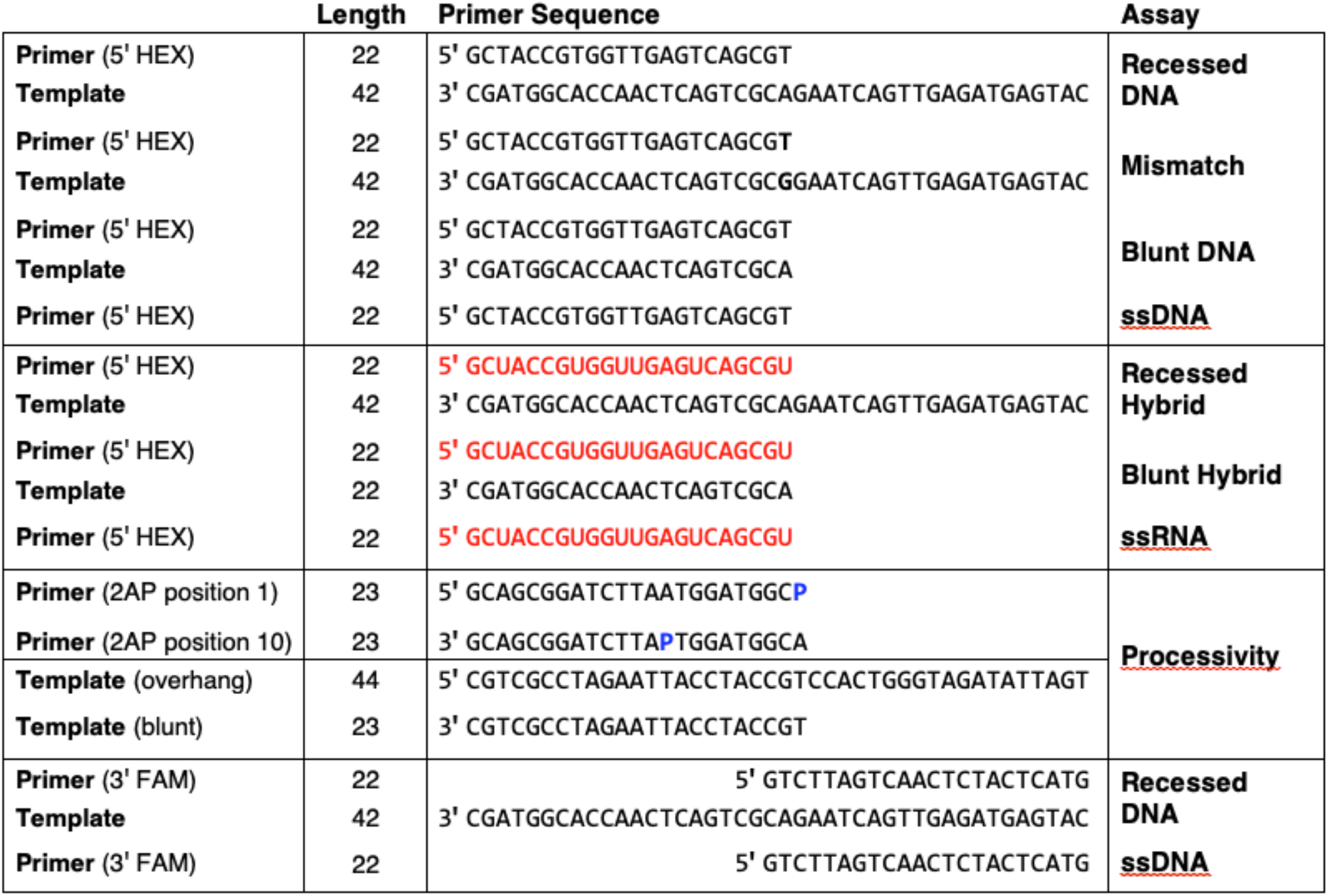
Annealed substrates for exonuclease and primer extension assays. Primer sequences displayed 5’-3’ and template sequences are displayed 3’-5’. Primers contain a 5’ hexachlorofluorescein (HEX) label, a 5’ 2-aminopurine (2AP) or a 3’ fluorescein (6-FAM) label. Black lettering indicated DNA sequence, red letters indicate RNA sequence, **bold** indicates mismatched nucleotides, and **blue P** indicates 2AP at either the 1^st^ or 10^th^ position.

### DNA substrate preparation

Two separate reactions were prepared in annealing buffer (10 mM Tris-HCl, 50 mM NaCl, 1 mM EDTA, pH 7.8) One reaction contained unlabeled primer (22mer, RNA or DNA) and template (42mer) in a 1:1.25 ratio, while the other contained 5**’** HEX-labeled 22mer (RNA or DNA) and unlabeled 42mer in a 1:1.25 ratio. Both reactions were heated to 95°C for 5 min and cooled to room temperature before being combined in a ratio of 1:20 labeled:unlabeled substrate (8.4 μM).

### Sequence Alignments and de novo Modelling

Representative Kinetoplastid POLIB protein sequences were retrieved from tritrypdb.org (release 56, 15 Feb 2022) that is hosted through the Eukaryotic Pathogen, Vector and Host Informatics Resource (veupathdb.org) and representative family A DNA polymerase domain sequences were aligned using NCBI COBALT multiple alignment tool.^14,15,16^ Alignments were viewed in Snapgene (version 6.0.2). Structure based alignment of TbPOLIB with *E. coli* Klenow (PDBid = kln1) defined residues 414-1400 as the POLA domain and were subsequently used for the design of truncated TbPOLIB for recombinant protein expression. DEDD family exonuclease motif sequences were aligned using the NCBI COBALT. De novo modeling of TbPOLIB (414-1400) was accomplished using the standard RoseTTAfold submission accessed via the RosettaCommons web-interface Robetta server on March 21^st^, 2022 (https://robetta.bakerlab.org/).^17,18^

### Plasmid Preparation for Bacterial Expression

TbPOLIB (UniProt Accession Q8MWB4) coding region (residues 414-1400) was codon optimized for expression in *E. coli,* synthesized and cloned into the pET100/D-TOPO vector (Geneart, Thermofisher) (Table S1A), resulting in the addition of an N-terminal 6x His-tag. This truncated version of TbPOLIB is termed IBWT for this study. Quikchange Lightning Multi Site-directed mutagenesis kit (Agilent Technologies) was used to introduce catalytic site point mutations (Table S1B) to IBWT resulting in IBPol- and IBExo-variants. Mutations were verified by sequencing, and all truncated POLIB variants were transformed into BL21(DE3) chemically competent cells (Invitrogen) for protein expression.

### Protein Purification

Purification scheme was identical for the three POLIB variants. Four liters of cells were grown in Luria Broth with shaking at 37°C until OD600 reached 0.4-0.8. Protein expression was induced with 40 μM final concentration IPTG and incubated 16-18 hours at room temperature (~22°C) with shaking. Cells were harvested by centrifugation, washed with phosphate buffered saline (PBS, 137 mM NaCl, 2.683 mM KCl, 10.144 Na_2_HPO_4_, 1.764 mM KH_2_PO_4_, pH 7.4) and stored at −20°C. Frozen pellets (8.7-11.6 g) were resuspended in Lysis Buffer (10 mM NaH_2_PO_4_, 10 mM Na_2_HPO_4_, 500 mM NaCl, 10 mM imidazole, pH 8.0) and lysed by passing 2-3 times through a microfluidizer (Microfluidics). After lysis, all steps were performed at 4°C. Lysate was centrifuged at 17090 x g for 1 hour, then the cleared bacterial lysate was passed through a 0.22 μm PES filter (Genessee Scientific) and then applied to a 5 mL packed Ni-NTA agarose beads (Goldbio) equilibrated with Lysis buffer. The beads were washed with 100 mL lysis buffer followed by 50 mL of Wash Buffer (10 mM NaH_2_PO_4_, 10 mM Na_2_HPO_4_, 500 mM NaCl, 40 mM imidazole, pH 8.0). Protein was eluted using 20 mL Elution Buffer (10 mM NaH2PO4, 10 mM Na_2_HPO_4_, 20 mM NaCl, 500 mM imidazole, pH 8.0) diluted 10-fold in Q Buffer (20 mM Tris, 1 mM DTT, pH 8.0) and applied to a pre-equilibrated (Q Buffer) 5 mL HiTrap Q-column (General Electric) using an ÄKTA pure chromatography system. Protein was eluted with a 0-500 mM NaCl linear gradient in Q buffer (75 ml). 1.5 mL fractions were collected. Fractions containing POLIB were identified by SDS-PAGE followed by staining with Coomassie blue. Fractions with active enzyme were pooled, concentrated (5 μM - 35 μM), and buffer exchanged into storage buffer (20 mM Tris, pH 8.0, 100 mM NaCl, 1 mM DTT, 20% glycerol) using Millipore Amicon Ultra Centrifugal Filter Units (50 kDa cutoff).

Protein concentration was determined using A280 (Nanodrop, Thermofisher), the molecular weight and the extinction coefficients of the POLIB variants predicted by Expasy ProtParam (https://web.expasy.org/protparam/). Small aliquots (10-20 μL) of protein were flash frozen in liquid nitrogen and stored at −80°C. Replicates of all assays were performed with three different preparations of the relevant POLIB variant.

### Gel-Based Exonuclease Activity Assays

The incubation mixture contained, in a final volume of 50 μL, 50 mM Tris-HCl (pH 7.0), 1 mM DTT, 0.1% BSA. As substrate, 840 nM of single-stranded (ssDNA) or 840 nM of doublestranded DNA (dsDNA) was used (Table 1). The amount of each POLIB variant (25 nM or 50 nM) was adjusted to obtain data points for initial reaction rates, as noted.

Reactions were initiated with the addition of 5 mM metal ion (Mg^2+^ or Mn^2+^) and incubated at 37°C for the indicated times. Time points of reactions were taken by removing 10 μL and quenching by adding an equal volume of quench buffer (0.1 M EDTA, 80% formamide, 0.1% Orange G) at designated time points. Quenched reactions were heated to 95°C for 10 min. Reactions were analyzed by electrophoresis in 7.5 M urea/20% polyacrylamide gels, visualized on a Typhoon™ FLA 9500 imager (GE) and band intensity was determined using Image J.^19^ Percent of 22mer remaining in exonuclease assays was calculated using Equation 1:

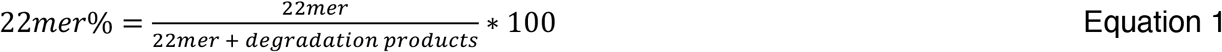

where *22mer* represents the total signal coming from the 22mer and *degradation products* represent the total signal coming from all the bands below the 22mer substrate.

Exonuclease rates were calculated using Equation 2:

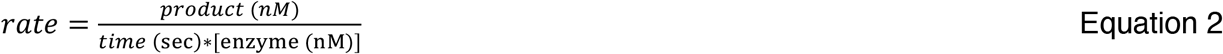

where *product* represents the concentration of all product bands in nM, *time* represents the time point at which the reaction was quenched and *enzyme* represents the concentration of enzyme in nM.

### Gel-Based Primer Extension Assays

Standard exonuclease reaction mixtures (50 μL) were used with the addition of 500 μM nucleotides (dNTP or rNTP) and increased POLIB variant concentration (200 nM). Immunoprecipitation of epitope-tagged Pol I-like paralogs from *T. brucei* cell extracts were performed at previously described.^10^ To assay immunoprecipitated protein for activity, 5 μL protein bound to IgG beads was added to each reaction, and the whole reaction was quenched after 60 minutes with the addition of 50 μL quench buffer. Quenched reactions were heated to 95°C for 10 min. Reactions were analyzed by electrophoresis and ImageJ using the same protocols as exonuclease Urea-PAGE assays. Percent extension product was calculated using Equation 3:

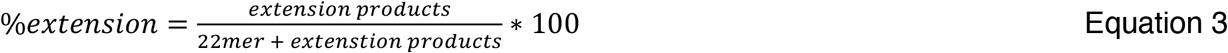

where *extension products* is the total signal from all the bands above the 22mer and *22mer* represents the signal from the 22mer substrate.

### 2-Aminopurine Exonuclease Processivity Assays

Each reaction (25 μL) consisted of 5 μM DNA substrate (Table 1), 50 mM Tris, 1 mM DTT, 0.1% BSA at pH 7.0. Reactions were preincubated at 37°C for 5 minutes before initiating with the addition of IBWT and were analyzed in a black 384 well plate. Relative fluorescence was measured using a SpectraMax M2e Microplate reader (excitation wavelength 310 nm and emission wavelength 375 nm) every 10 seconds. Free 2AP was calculated using a standard curve of 2AP (Alfa Aesar) measured using the same conditions as the processivity reactions. Each reaction was run in triplicate. Dynafit was used for global fitting of the full progress curves to determine the ratio of the lower or upper boundaries of the rate constants for 1^st^ position 2AP substrates and 10^th^ position 2AP substrates, the two scripts used to fit the data can be found in Supplementary Figures 1&2.

### Trypanosoma brucei DNA Constructs

POLIA was amplified from gDNA from TREU927 cells using primers POLIA F and POLIA R (Table S2) and cloned into pLEW79-MHTAP using the BamHI site. The protein encoding sequence of POLIB was amplified from *T. brucei* procyclic form Lister 427 gDNA using primers POLIB F and POLIB R adding XhoI and XbaI cut sites (Table S2) and ligated into pLew100-PTP^Puro^ digested with the same restriction enzymes, generating pLew100-FLPOLIB-PTP^Puro^.^19,10^ A POLIB gene fragment (1-1053 bp) was obtained from Genescript in pUC57 to replace the natural corresponding POLIB sequence in pLew100-FLPOLIB-PTP^Puro^. The gene fragment included a recoded region (249-801 bp) to use in unrelated RNAi complementation experiments. The gene fragment was amplified with primers POLIB Rc F and POLIB Rc R (Table S2) for subsequent Gibson assembly with pLew100-FLPOLIB-PTP^Puro^ digested with XhoI and BamHI. Resulting vector and gene fragment were combined via Gibson Assembly (Takara In-fusion Kit) generating pLew100-FLPOLIB_Rc_-PTP^Puro^. Polymerase and exonuclease catalytic site mutations were separately introduced to this construct using mutagenic primers POLIB M1, POLIB M2, and POLIB M3 (Table S2) and the QuikChange Lightning Multi-Site Mutagenesis Kit (Agilent) according to manufacturer’s protocol.

The *POLID* open reading frame (4887 bp), excluding the stop codon, was PCR-amplified from *T. brucei* 927 genomic DNA using primers POLID F and POLID R (Table S2). The PCR product was cloned into pCR-Blunt-II-TOPO (Invitrogen) to generate pTOPO-FLID-WT. Site-directed mutagenesis of *POLID* was performed on pTOPO-FLID-WT using the QuikChange Multi Site-Directed Mutagenesis Kit (Agilent) with primers POLID M1 and POLID M2 to generate pTOPO-FLID-Poldead. Due to the relatively large plasmid size of pTOPO-FLID-WT, non-mutagenic primers POLID M3 and POLID M4 (Table S2) which anneal to the pCR-Blunt-II-TOPO backbone, were also included in each mutagenesis reaction. TbPOLID variants wild type and polymerase-dead were cloned into pLew100-PTP^Puro^ via Mfel and BamHI sites by GENEWIZ, Inc. generating pLew100ID-WT-PTP^Puro^ and pLew100ID-poldead-PTP^Puro^.

### Trypanosoma brucei Culture and Transfection

Procyclic *Trypanosoma brucei brucei* 29-13 cells expressing T7 RNA polymerase and tetracycline repressor were maintained at 27°C in SDM-79 medium supplemented with heat-inactivated fetal bovine serum (15%), G418 (15 μg/ml), and hygromycin (50 μg/ml).^21^ Transfected cell lines were additionally supplemented with the appropriate selectable drug and the resulting transgenic variants used in this study are listed in Table S2B.

Generation of overexpressing POLIC variant cell lines was previously described.^10^ 10 μg pLEW79IA-MHTAP was NotI linearized and transfected into 29-13 procyclic cells by electroporation.^8^ 7.5 μg pLew100-PTP^Puro^ plasmids encoding full length POLIB and POLID variants were NotI linearized and transfected into 29-13 cells via Amaxa Nucleofection (Parasite Kit, Lonza). Following selection with 1 μg/mL puromycin for pLew100 based constructs and 2.5 μg/mL phleomycin for cells transfected with pLEW79IA-MHTAP, clonal cell lines were obtained via limiting dilution as previously described.^8^

Overexpression of all variants was induced by addition of 1 μg/mL tetracycline to the cultures and supplemented with 0.5 μg/mL on days dilutions were not performed. Clones were selected for use based on highest expression of inducible protein. Cultures were induced for 48 hours before cells were collected for immunoprecipitation.

### Western Blot and SYPRO™ Ruby Protein Detection

Proteins were separated via SDS-PAGE on 8% polyacrylamide gels, followed by 16-hour transfer onto a PVDF membrane. Membranes were blocked in Tris-buffered saline (TBS) containing 5% non-fat dry milk. PTP-tagged or MHTAP-tagged protein variants were detected using the peroxidase-anti-peroxidase soluble complex (PAP) (1:2000, Sigma) as previously described.^10^ Detection of 6x His-tagged recombinant POLIB variants was performed with primary antibody Penta•His (1:1000, Qiagen) for one hour, and subsequently incubated with secondary rabbit anti-mouse IgG horseradish peroxidase conjugate (1:4000, Zymed) for one hour. Signal was detected using Pierce™ ECL Western Blotting Substrate with an Amersham ImageQuant 800 (Cytiva). Sypro^TM^ Ruby (Thermofisher) detection of total protein was performed following SDS-PAGE according to manufacturer’s protocol.

## RESULTS

Kinetoplastid organisms are the only group of eukaryotes that possess four Family A DNA polymerase paralogs exclusively targeted to the mitochondrion.^5,7^ Among these paralogs, the Family A Pol domains (PolA) share only 25-35% amino acid identity, suggesting specialized roles for each.^5^ In addition to the conserved C-terminal PolA domain, only the POLIB and POLID paralogs contain predicted exonuclease (exo) domains. Although the POLID exonuclease domain shares similarity with those of other Family A pols, the POLIB exo domain is markedly different in sequence and position.

### De novo Protein Structure Prediction of POLIB Reveals a Large Insertion in the Thumb Subdomain

Predictions of POLIB domains using the SuperFamily database show two stretches of amino acids that have homology to Family A polymerases (414-660 and 1031-1400 aa). (Figure 1A). ^22^ Embedded within the PolA domain is a region with homology to the DnaQ 3’-5’ exo superfamily (763-956 aa). The first 30 aa are predicted to be a mitochondrial targeting sequence (TargetP-2.0)^23^ but the remainder of the N-terminal region (through aa 413) does not have homology to any characterized proteins and is referred to as the uncharacterized region (UCR). Attempts to gain structural insight into the unusual POLIB domain arrangement using homology modeling of amino acids 414-1400 with the *E. coli* Pol I structure revealed a canonical right-hand Pol domain structure (fingers, palm, thumb subdomains) with a large unstructured insertion in the thumb domain that spanned the C-terminal amino acids 631-1030. When modeled separately, the insertion had homology to *E. coli* RNaseT, a DnaQ exo superfamily member.

**Figure 1:**
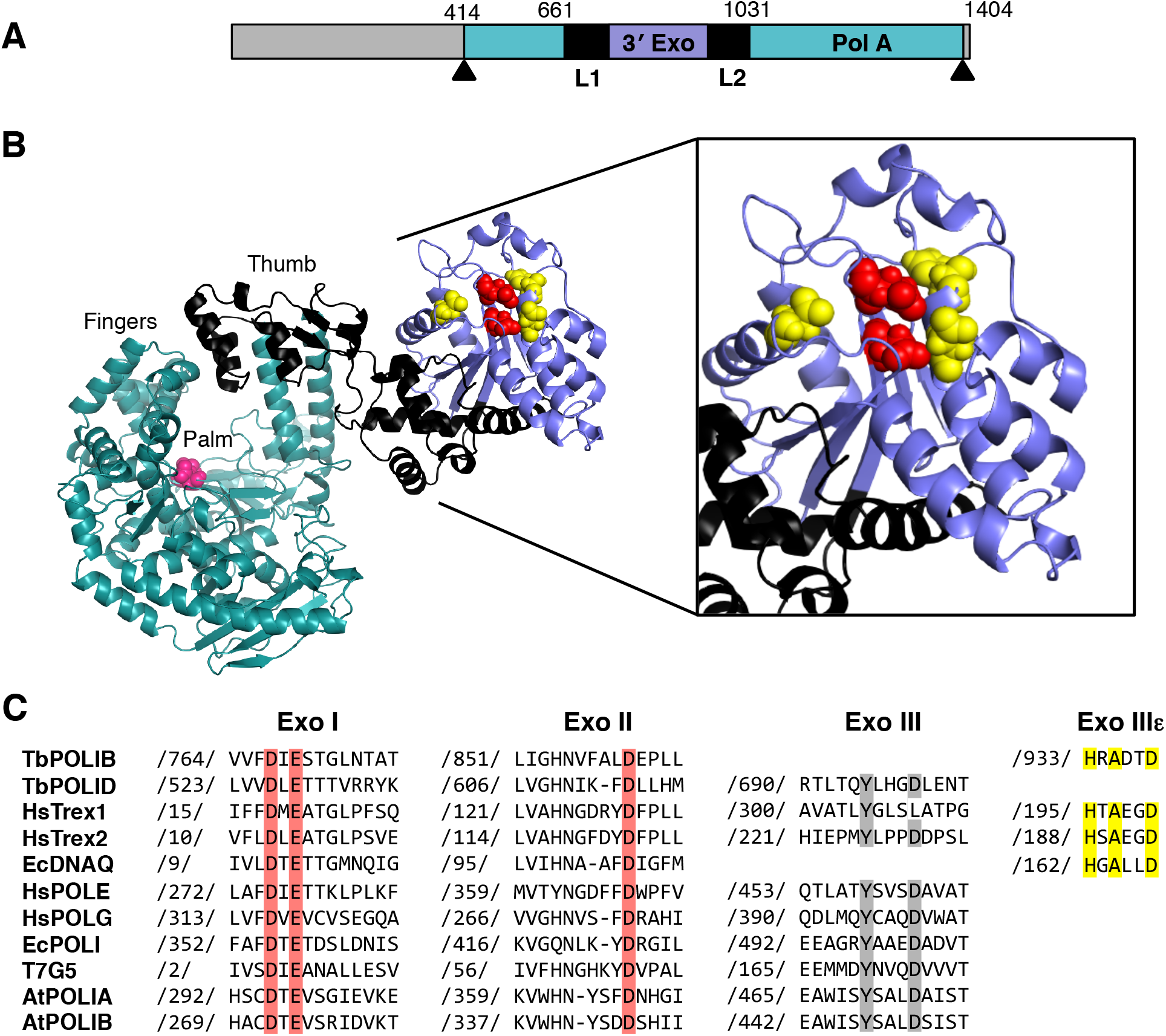
Divergent Structure of TbPOLIB. (A) Schematic representation of TbPOLIB with predicted polymerase (Pol) and exonuclease (Exo) domains. Mitochondrial targeting sequence, 1-30; N-terminal uncharacterized region, 31–413; Linker region 1 (L1), 661–761; Exo, 762–955; Linker region 2 (L2), 956– 1030 and pol domain, 413–660 and 1031–1364. Arrowheads mark the region used for de novo modeling (414-1400). (B) RoseTTAfold model of TbPOLIB. Teal, pol domain shows as canonical “right-hand’’ configuration with fingers, palm and thumb, fingers subdomains; purple, thumb insertion with homology to exonuclease; black, linker regions. Magenta, Pol domain active sites (only one active site residue is seen in this orientation); Red, Exo domain Asp767 and Glu769; Yellow, additional residues associated with DEDDh exonucleases. (C) Alignment of conserved Exo motifs for selected DEDD 3’-5’ exonucleases. Red, three active site carboxylates; Yellow, DEDDh consensus motif (HxAxxD); Grey, DEDDy consensus motif (YxxxD).

Recent developments in *de novo* protein structure prediction using deep learning now allow for predicted protein structures with atomic accuracy (Baek 2021, Jumper 2021).^17,24^ We used the *de novo* modelling platform RoseTTAfold to gain insight into how the two POLIB annotated enzymatic domains would fold as part of the same structure. Figure 1B depicts a model with a local Distance Difference Test (lDDT) confidence score of 0.75, indicating a good quality model.^25^ In the plot of predicted error of position of the residues in the model, the majority falls under 5 ångströms with the main exception being the C terminus (Figure S3A). The Pol domain right-hand structure contains the previously identified invariant aspartic acids (D1117 and D1309) essential for nucleotidyl incorporation in the palm subdomain as expected. An amino acid alignment of various PolA domains highlights the conserved Motifs A, B and C and a key difference for POLIB, which is the presence of HDS instead of the invariant HDE found in Motif C in all other Family A pols insertion (Figure S4).

The RoseTTAfold model depicts the large 369 aa insertion as a projection from the tip of the thumb. Residues 763-956 fold into a separate domain that is bounded by flexible linker regions (L1 and L2). Smaller thumb domain insertions have been described in other Family A Pols that contribute to increased processivity (T7 Pol, human Pol θ) or translesion DNA synthesis *(A. thaliana* PolIA).^11,12,26^ Additionally, proofreading Family A members have the exo domain located under the palm subdomain (*E. coli* Pol I, Pol γ).^27,28^ Thus far, POLIB is divergent from all other Family A proteins containing a unique substitution within Motif C, and the largest thumb insertion with similarity to an exo domain. Interestingly, these divergent features are conserved among all the POLIB sequences from Kinetoplastid organisms sequenced to date (Figure S5).

A similar conformation was predicted for the full-length POLIB model retrieved from the Alphafold database depicted with (Fig S3B) and without the UCR present in the model (Figure S3C). The position of the UCR relative to the other domains has low confidence based on the predicted aligned error so its position relative to the rest of the protein is unreliable in this model (Figure S3D).

The DnaQ exo superfamily contains three conserved motifs Exo I, Exo II and Exo III that are clustered around the active site and participate in a two-metal ion catalysis. This superfamily is also called the DEDD based on four negatively charged invariant residues present in the conserved motifs and can be further classified into DEDDy or DEDDh proteins depending on the variation found in motif III (YxxxD or HxAxxD, respectively).^29^ POLIB contains the essential active site residues characteristic of the DEDDh subfamily of exos (Figure 1B, C). DEDDh exos includes *E. coli* DnaQ, the proofreading subunit for *E. coli* Pol III, and human TREX1 and TREX2 which non-processively degrade cytosolic ssDNA and dsDNA.^30^ The 3’-5’ exo domain of POLID, a paralog of POLIB, is more similar to the DEDDy proofreading domains of human Pol g and Pol ε.

### POLIB Thumb Subdomain Insertion is an Active 3’-5’ Exonuclease

To evaluate whether the POLIB insertion was an active 3’-5’ exo, we generated codon optimized His-tagged recombinant POLIB variants for expression and purification from *E. coli.* The variants were truncated to eliminate the UCR and include IBWT, IBExo- (D767N, E769Q), and IBPol-, (D1117A, D1309A). All variants were purified to near homogeneity with the expected molecular weight of 115.6 kDA using Ni affinity and anion exchange chromatography. SYPRO ruby staining demonstrated that the purity of the POLIB variants was higher than 96% (Figure S6). Typical yields were 0.575-1.625 mg of pure protein per liter of induced culture.

The 3’-5’ exonuclease activity of POLIB variants was measured using a 5’ hexachlorofluorescein (HEX)-labeled 22mer primer annealed to an unlabeled 42mer template in the absence of dNTPs. Degradation products were visualized on a Urea-PAGE gel. IBWT exhibits 3’-5’ exo activity that is completely ablated in the IBExo-variant, while IBPol-displayed reduced degradation activity on a recessed dsDNA template (Figure 2A). IBWT degraded 15% of the primer to 21 mer or smaller products within the first minute of the reaction, whereas IBPol-only degraded 5% during the same time. At longer time points, from 2 to 8 min, intermediate degradation products accumulated but products smaller than 10mer were only observed for IBWT.

**Figure 2.**
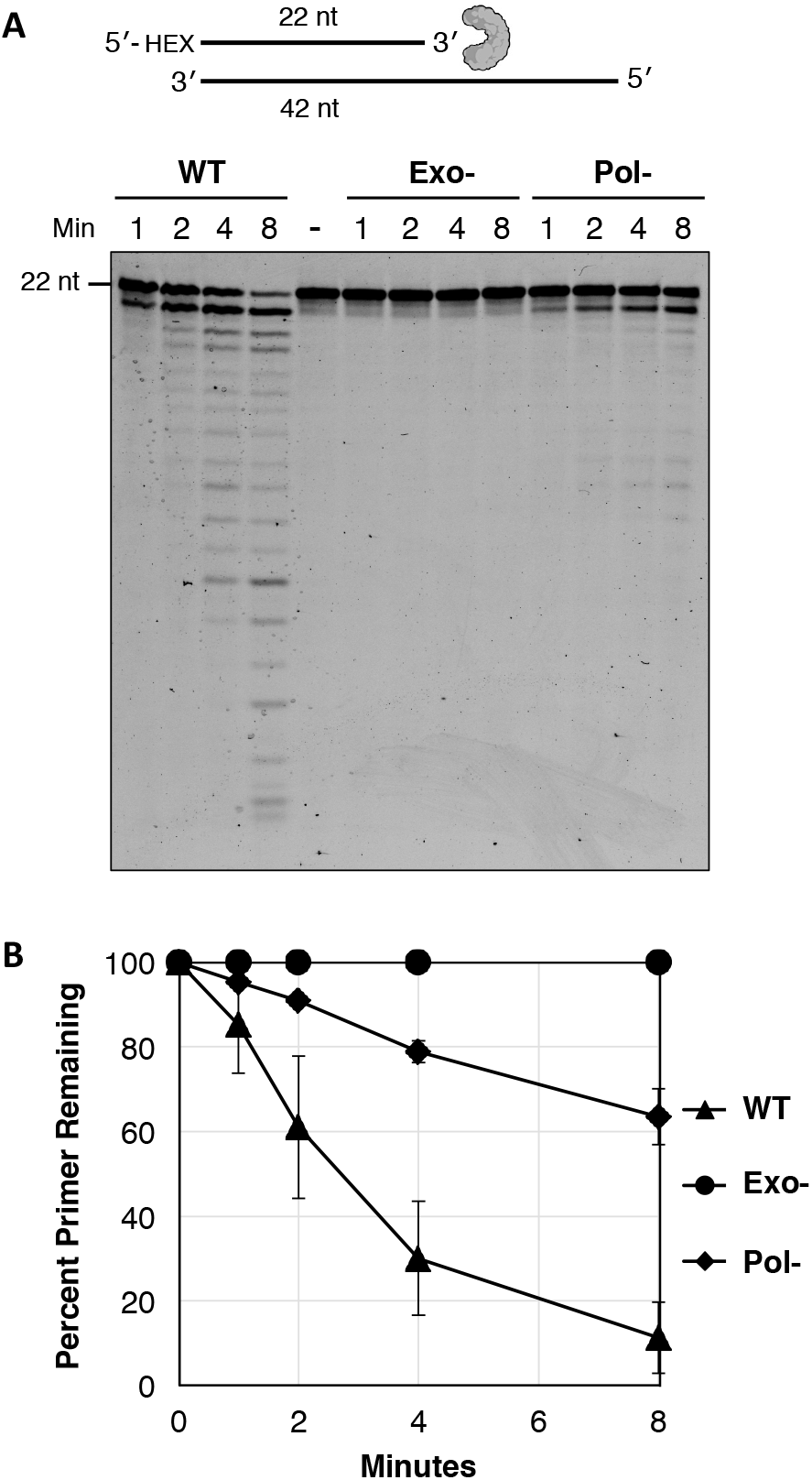
Exonuclease activity of POLIB variants. (A) Examination of 3’-5’ exonuclease activity using 840 nM recessed 5’ HEX-labeled primer/template dsDNA substrate, 50 nM POLIB of each variant in the absence of dNTPs. Reactions stopped at the indicated times to evaluate progression of reactions. Representative image from multiple experiments. No protein control Time=0, N. (B) Quantification of dsDNA degradation of POLIB variants. N=3, error bars represent standard deviation.

Quantification of primer degradation for the POLIB variants demonstrated that catalytic mutations in IBPol-impacted the rate of DNA degradation about 2-fold, suggesting that there may be crosstalk between the Pol and Exo domains of POLIB, a characteristic described in other well-characterized DNA polymerases such as human Pol g.^28^ No 5’-3’ exonuclease activity was detected when using 3’ fluorescein (6-FAM)-labeled single stranded or double stranded DNA substrates compared to the T5 exo control (Figure S7).

### Substrate Preference for POLIB Exonuclease Activity

Gel-based assays using the HEX-labeled DNA and RNA substrates were also used to determine substrate preference for IBWT exo activity (Figure S8). Some proofreading DNA polymerases contain an additional 3’-5’ exonuclease activity that can be used to remove an incorrectly added nucleotide during DNA extension. Proofreading domains typically display a preference for removing mismatched nucleotides over correctly paired matches, including those domains in human Pol γ and *E. coli* Pol I.^31,32^ We tested whether POLIB displays a preference for mismatch removal by observing the relative rate of removal of a mismatched terminal nucleotide versus a correctly terminal nucleotide on a recessed 3’ DNA substrate (Figure 3A). No significant difference was observed in the relative rates of these reactions, suggesting POLIB does not have a preference for mismatched substrates.

**Figure 3.**
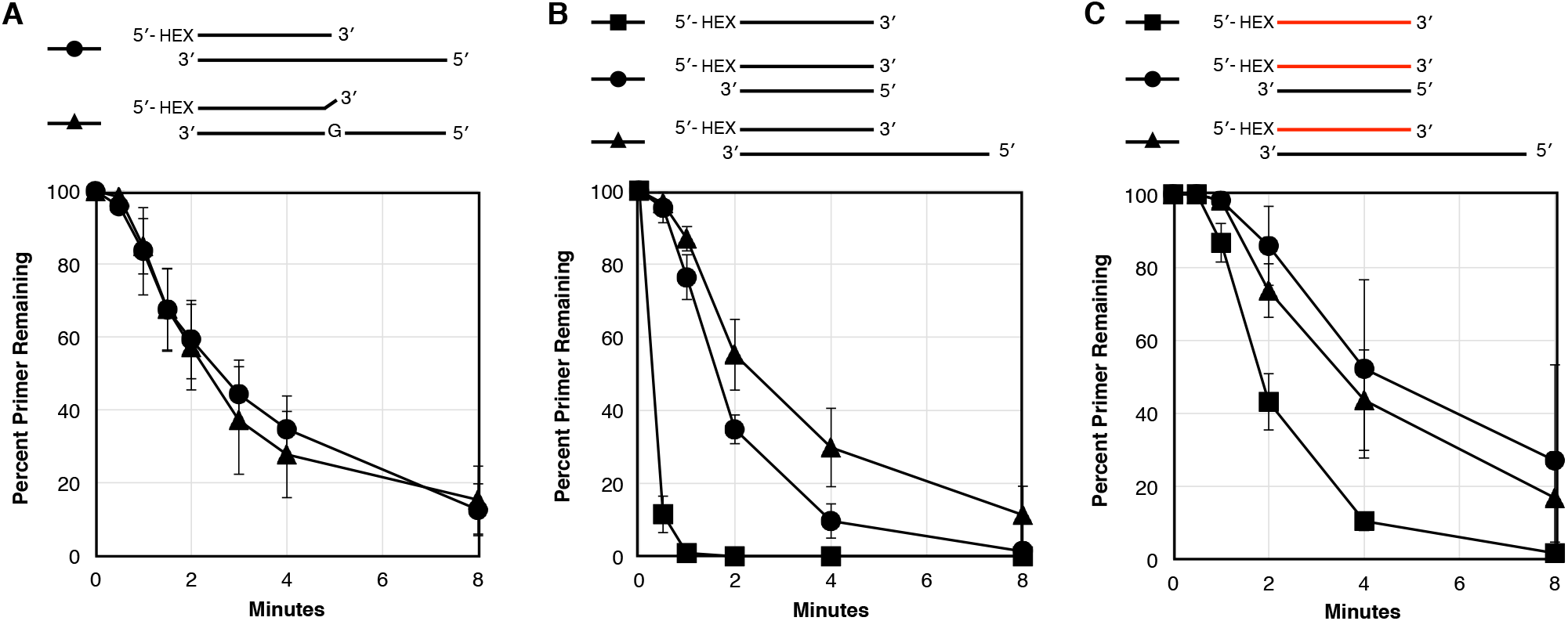
POLIB exonuclease template preference. Quantification of gel-based exonuclease assays. Standard reactions contained 25 nM IBWT for single stranded substrates or 50 nM for double stranded substrates in order to observe the linear range of these reactions. Reaction time points were run on Urea-PAGE gels and quantified using image J. Red oligo: RNA, Black oligo: DNA. (A) Relative rates of primer degradation with either 3’ terminal matched or mismatched basepairs. (B) Relative rates of degradation of DNA oligos. ssDNA reaction used 25 nM POLIB and 840 nM DNA, dsDNA reactions used 50 nM POLIB and 840 nM DNA. (C) Relative rates of degradation of 840 nM ssRNA with 25 nM POLIB, or RNA in an RNA-DNA hybrid with 50 nM POLIB.

We also assayed the exonuclease activity on different DNA templates to determine what may be the preferred substrates of POLIB. Due to the higher rate of degradation of POLIB on single-stranded oligos, IBWT concentration was reduced to 25 nM in reactions with these substrates. IBWT successfully degraded DNA from the 3’ end and was capable of degrading both ssDNA and dsDNA with a 5’ overhang or blunt dsDNA. IBWT degraded ssDNA at a faster rate than either dsDNA substrate, with an average rate of 0.9914 ± 0.0560 s^-1^ for ssDNA, compared to 0.0626 ± 0.0136 s^-1^degradation of the recessed DNA substrate and 0.09133 ± 0.0055 s^-1^ for blunt dsDNA (Figure 3B). On average, IBWT degraded blunt dsDNA more rapidly, and appeared to remove the terminal nucleotide of the blunt DNA substrate more rapidly than the terminal nucleotide of the recessed DNA substrate. However, POLIB degraded subsequent nucleotides more rapidly in the recessed DNA substrate than the blunt substrate, as seen by the increase of shorter degradation products. (Figure S8). This could be due to an affinity of IBWT to longer DNA templates. POLIB is also capable of degrading ssRNA and RNA/DNA hybrids, although at lower rates than DNA degradation (Figure 3C). No detectable degradation occurred in the first 30 seconds of the ssRNA degradation reaction, suggesting IBWT has a strong preference for degrading ssDNA. On average, the rate of degradation for the ssRNA oligo was 0.1591 ± 0.0216 s^-1^, as compared to a rate of 0.0369 ± 0.0103 for the recessed RNA and 0.0198 ± 0.0151 for the blunt RNA/DNA hybrid.

### POLIB degrades DNA substrates in a Non-processive Manner

DNA substrates containing 2-aminopurine (2AP) at either the 3’ end (1st position) or at the 10^th^ internal nucleotide (10^th^ position) were used to evaluate the processivity of IBWT on three different substrates: ssDNA, recessed dsDNA, or blunt dsDNA. Single-hit conditions were used, where the concentration of the DNA substrate was much higher than IBWT (25x higher for dsDNA substrates, 200x higher for ssDNA substrates).^33^ For all the three substrates, a significant delay between the removal of the 1^st^ position 2AP and the 10^th^ position 2AP was observed. The lag period that is observed for the 10th position DNA substrate indicates that multiple binding and nucleotide excision events must occur prior to the enzyme reaching the 10th position (Figure 4).

**Figure 4.**
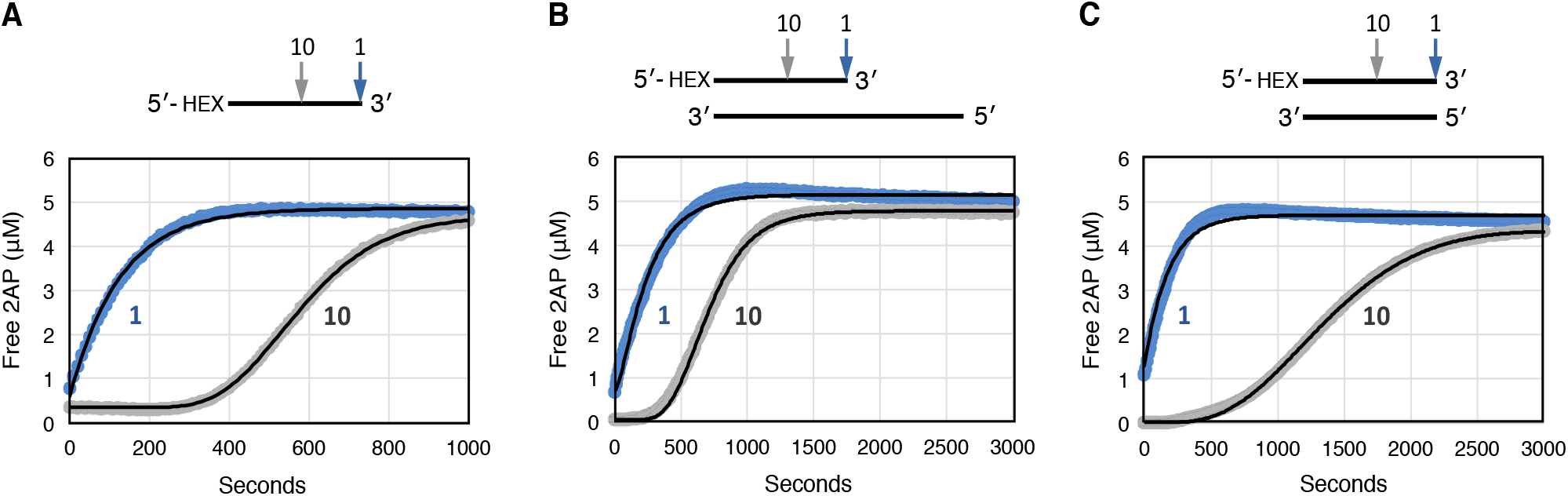
Exonuclease Processivity of POLIB. Standard reactions contained 5 μM of DNA substrate with 2AP either at the 1^st^ position (1AP) or the 10^th^ position (10AP). Progress curves were fitted using Dynafit to determine upper or lower boundaries of *k*_exo_ and *k*_off_. Blue circles, 1AP data points; grey circles, 10AP data points; black line, fitted progress curve. (A) Representative progression curves for ssDNA substrate degradation by 50 nM IBWT. (B) Representative progression of reactions for recessed dsDNA substrate degradation by 200 nM IBWT. (C) Representative progression curves for blunt dsDNA substrate degradation by 200 nM IBWT.

The lower or upper limit of the *k*_exo_ and *k*_off_ values were determined by global fitting of the progress curves for the 1st and 10th 2AP positions with Dynafit (Figures S1, S2). This ratio of these two rate constants can be used to determine the likelihood of whether POLIB will dissociate from a DNA substrate or remain bound to excise another nucleotide. This ratio was 95.92 ± 8.9 for ssDNA, indicating very low processivity. For the recessed dsDNA substrate, the ratio of the upper limit of *k*_exo_ and *k*_off_ is 28.949 ± 30.988, also demonstrating low processivity of IBWT (Figure 4B). For blunt dsDNA, this ratio is 2.429 ± 0.462, which also indicates POLIB is more likely to dissociate from this substrate after removing a single nucleotide than removing another in a single binding event (Figure 4C).

For ssDNA substrates, the excision of the 1 ^st^ position 2AP occurred at a slower rate than subsequent nucleotides, which could indicate a lower affinity for the 2AP nucleotide or the presence of a slow activation step prior to the onset of activity (i.e., enzyme hysteresis) (Figure 4A). This is also the case for blunt dsDNA but was not observed in the case of the recessed dsDNA substrate. This possibly indicates a reduced affinity of IBWT to the shorter products of degradation from the blunt 22mer in comparison to the longer 42mer template of the recessed substrate. These substrate-dependent changes in rate as the enzyme proceeds further into the substrate were also observed on gel-based exo assays, as noted (Figure 3).

### pH Impacts Competition Between DNA Extension and Degradation Activities of POLIB

Two other pols involved in kDNA maintenance, Pol β and Pol β-PAK, incorporate nucleotides most efficiently at pH 9.^4^ We therefore predicted POLIB would catalyze more efficiently at a basic pH. Notably, we observed a change in IBWT preference between degradation and extension activities that was dependent upon pH during characterization of recombinant POLIB (Figure 5A).

**Figure 5.**
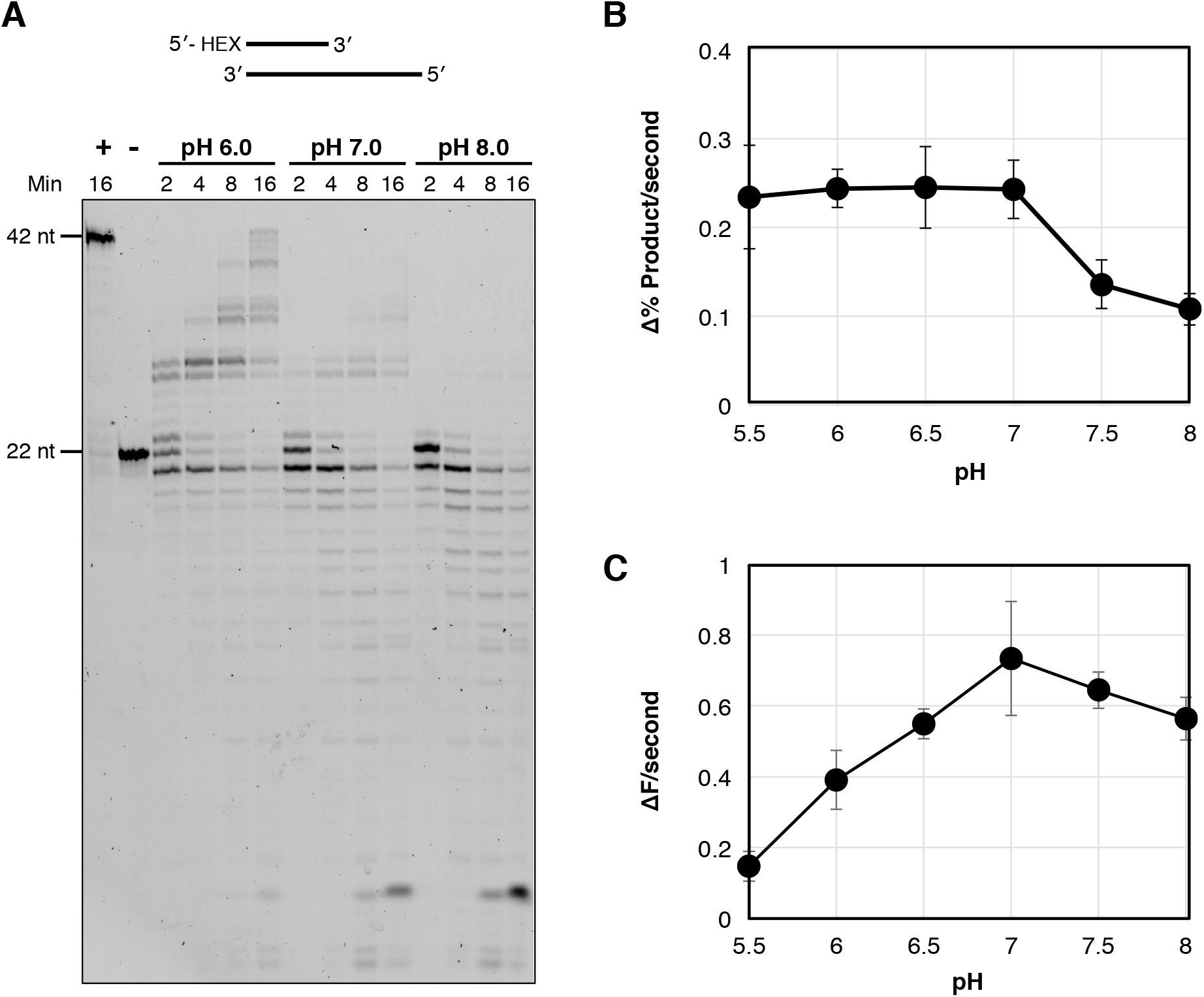
Impact of pH on POLIB exonuclease and extension activities. A) Representative image of IBWT extension and degradation products assayed using standard reaction buffer with varying pH conditions and 500 μM dNTPs. 1U Klenow, +; Time=0, pH 7.0, N. (B) Rates of extension determined using a gel-based assay. Standard reaction buffer was used with varying pH conditions, 200 nM IBExo-840 nM oligo substrate, and 500 μM dNTPs. Rate determined using quantification of original primer and extension products. N=3, error bars represent standard deviation. (C) Rates of 2AP excision from recessed dsDNA substrate varying pH conditions, 200 nM IBWT and 5 μM substrate in each reaction.

IBWT was used to demonstrate the relative rate of extension and degradation of a 5’ Hex-labeled recessed DNA template at pH conditions of 6.0, 7.0, and 8.0. At pH 8.0, minimal nucleotidyl incorporation was observed but shorter exonuclease products build up through the course of the reaction. At pH 7.0, both extension products and degradation products are being formed through the course of the reaction. At pH 6.0, more extension products are formed than degradation products (Figure 5A). We hypothesized that this pattern was due to separate pH optimums for the nucleotidyl incorporation and exonuclease activities of POLIB. To test this, we used IBExo- (lacking competing exonuclease activity) to observe the rate of extension in varying pH conditions using the gel-based assay. POLIB extension activity was relatively similar from pH 5.5 – 7.0 but decreased in pH 7.5 and 8.0 by about half (Figure 5B).

To test the rate of nucleotide excision across pH conditions, we used a primer with a terminal 3’ 2-aminopurine (2AP), an adenine analog that is quenched by base-stacking and fluoresces when released from the primer.^33^ Using a recessed dsDNA substrate, POLIB was able to degrade DNA most rapidly at a pH of 7.0. The rate decreased under more acidic pH conditions, demonstrating a 3-fold lower rate of excision at pH 5.5 compared to pH 7.0. However, at pH 8.0 exonuclease activity only declines 23% from the rate at pH 7.0 (Figure 5C). Similar patterns were seen for the change in efficiency of exonuclease activity across pH conditions for a blunt dsDNA oligo and ssDNA (Figure S9). These results demonstrate that the pH tolerance for the nucleotidyl incorporation and degradation activities of POLIB are distinct and impact the activity preference of the enzyme.

We also investigated other factors that could have an impact of the rate of primer extension by IBExo-. Increasing the salt concentration of either NaCl or KCl, from 0 mM to 50 mM reduced the extension activity of IBExo- (Figure S10). A concentration of 150 mM of either salt eliminated any detectable extension. Another variable we tested was an exchange from using Mg^2+^ as the divalent cation to Mn^2+^. These reactions demonstrated that IBExo- incorporates dCTP at a similar rate with either divalent cation but was more likely to misincorporate dCTP past the correct basepair on the template with Mn^2+^. Furthermore, with Mg^2+^ as a divalent cation IBExo- did not extend the DNA primer with CTP, but was able to incorporate CTP in the presence of Mn^2+^ (Figure S11 A, B).

### POLIB Exhibits Preference for Extension from RNA primers

The invariant Asp in Motif C that is conserved in other Family A pols is replaced with a Ser in the Kinetoplastid lineage (Figure S3). A serine is present in this motif in some RNA pols such as T3 bacteriophage RNA Pol and *Saccharomyces cerevisiae* mitochondrial RNA Pol.^34,35^ Based on the Ser substitution, we hypothesized that the pol domain of POLIB may have a role in RNA metabolism. We evaluated the three POLIB variants in 60-minute extension reactions for extension activity using either an RNA or DNA primer is the presence of deoxyribonucleotides or ribonucleotides (Figure 6A).

**Figure 6.**
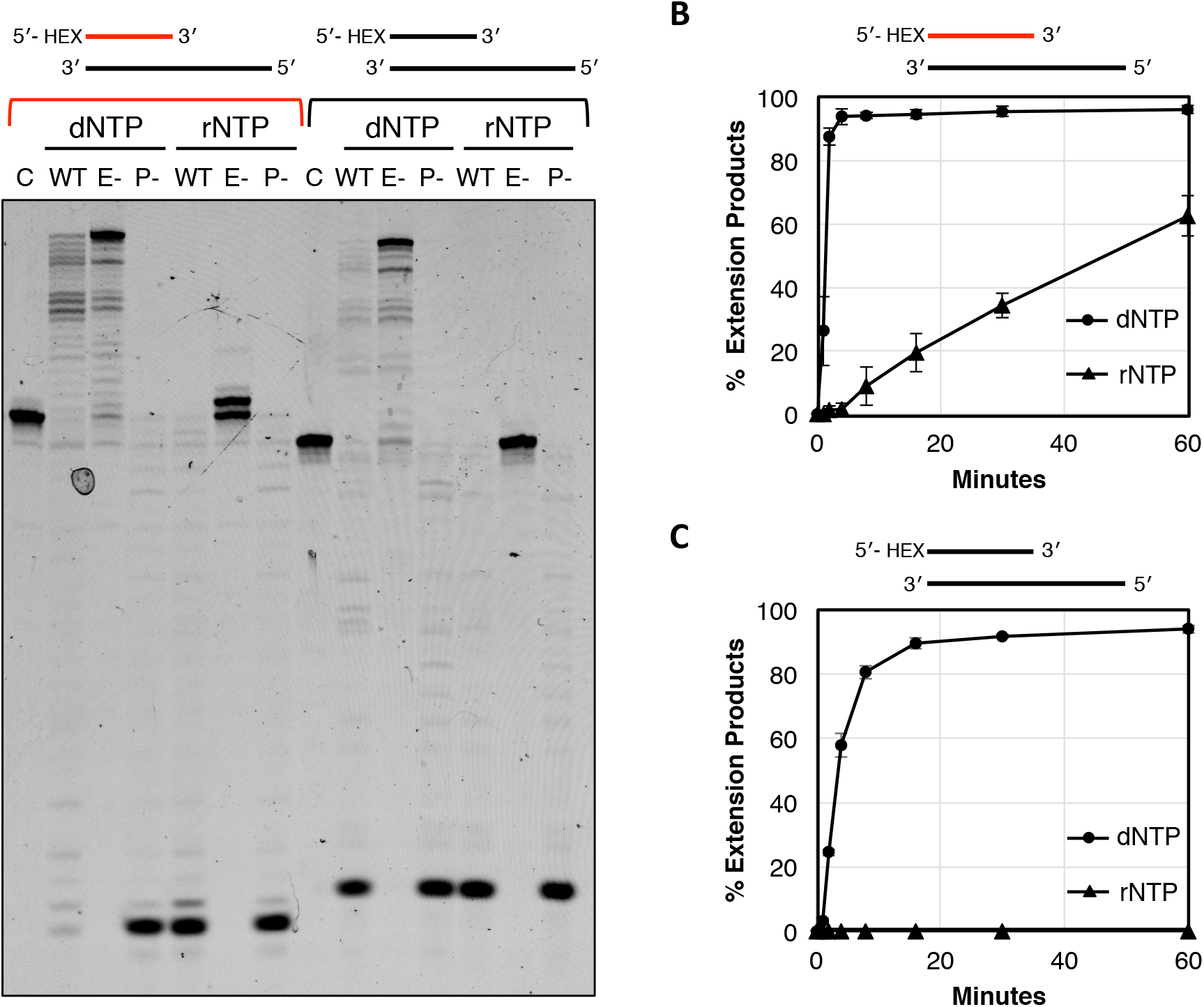
POLIB extension from RNA and DNA primers. Gel-based assays for primer extension, each reaction included 840 nM oligo, 200 nM POLIB variant, 500 μM dNTPs or rNTPs in standard assay buffer. Red oligo: RNA, Black oligo: DNA. (A) Representative gel-based assay for POLIB variant extension of DNA or RNA primers. and was quenched at 60 minutes. No protein control Time=0, C. (B) Quantification of gel-based assay time course of IBExo- extension from RNA primer, averages from 3 gels (C) Quantification of gel-based assay time course of IBExo- extension from DNA primer, averages from 3 gels, error bars represent standard deviation.

IBWT was capable of extending from an RNA primer utilizing dNTPs, and this extension activity was improved in IBExo- that lacks formation of degradation products and results in an increase in fully extended product (Figure 6A). When the reaction using IBWT included rNTPs instead of dNTPs, exonuclease activity outcompeted extension. IBExo- was capable of extending an RNA primer with rNTPS, although the majority of the products formed in this case are the result of a single nucleotide addition. Any extension of the RNA primer is ablated in reactions with IBPol-, although IBPol- is capable of degradation of both RNA and DNA primers.

With a template primed with DNA and the addition of dNTPs, IBWT formed more degradation products than with an RNA primer and fewer extension products. IBExo- produced more extension products than IBWT in these conditions as well. No IB variants were able to extend the DNA primer with rNTPs. In order to determine if primer composition impacts the rate of the POLIB nucleotidyl incorporation, we quantified the progress of extension reactions using Urea-PAGE over a 60-minute time course. IBExo- extended an RNA primer with dNTPs more efficiently than with rNTPs. (Figure 6B). Progress curves of IBExo- extension of a DNA primer with dNTPs were slower than extension from an RNA primer, with 3% of the DNA primer extended in the first minute of the reaction compared to 26% of the RNA primer (Figure 6B, C). IBExo- was not able to incorporate rNTPs from a DNA primer at all (Figure 6C). This primer-dependent activity may suggest a conformational change of the IB pol domain active site when bound to an RNA primer.

### Activity of Immunoprecipitated Mitochondrial POLI-like proteins

To evaluate whether truncation of the POLIB UCR impacts POLIB activities we immunoprecipitated POLIB variants and the other mitochondrial Pol I-like paralogs from *T. brucei* cell lines overexpressing full-length C-terminally PTP epitope-tagged proteins. Successful immunoprecipitation of variants was verified by SDS-PAGE and Western blot (Figure S12A). Equal volumes of PTP-tagged proteins still bound to IgG sephorase beads were assayed using standard buffer conditions, the HEX-labeled recessed DNA substrate. Initially pH 8.0 was used to mimic the predicted basic environment of the mitochondrial matrix.^36^

In addition to the Klenow control, POLIA, POLIC, and POLID wildtype proteins exhibited nucleotidyl incorporation while precipitates from untagged parental 29-13 *T. brucei* and beads only lacked any detectable activity. POLIC and POLID variants with alanine substitutions in the Pol domain conserved aspartic acid catalytic residues (ICpol- and IDpol-) ablate all detectable nucleotidyl incorporation activity (Figure 7A). The results for POLIC and POLICpol- agree with previously reported data.^10^ POLIBwt, POLIBpol-, and POLIDwt exhibited 3’-5’ exonuclease activity. POLIDpol- demonstrated significantly more degradation products than POLIDwt. Notably, POLIBwt exhibited only exonuclease activity even in the presence of dNTPs that is lost in the POLIBexo- variant. However, weak nucleotidyl incorporation was detected (Figure 7A). This suggests that the full-length POLIBwt exonuclease activity outcompeted nucleotidyl incorporation activity at pH 8.0. We next asked whether pH and primer composition might also impact extension activity of immunoprecipitated POLIB variants similar to that of truncated recombinant protein. Successful immunoprecipitations were verified by SDS-PAGE and Western blot (Figure S12B). POLIBwt and POLIBexo- immunoprecipitate was assayed under standard primer extensions conditions, with the pH varied as noted. Activity on RNA or DNA primers was compared using either a HEX-labeled recessed DNA substrate or a HEX-labeled recessed RNA/DNA hybrid. When immunoprecipitated POLIBwt was assayed at pH 8.0, more degradation products were detected than at lower pH conditions for both the DNA and RNA substrates similar to the results for truncated IBWT (Figure 7B). Degradation activity was favored over primer extension on a DNA primed substrate, but extension from an RNA primer was favored for both the POLIBwt and POLIBexo- variants. Some degradation products are detected in reactions with POLIBexo- possibly due to incomplete ablation of exonuclease activity in the full-length protein or a contaminating exonuclease in the immunoprecipitated protein sample (Figure 7B).

**Figure 7.**
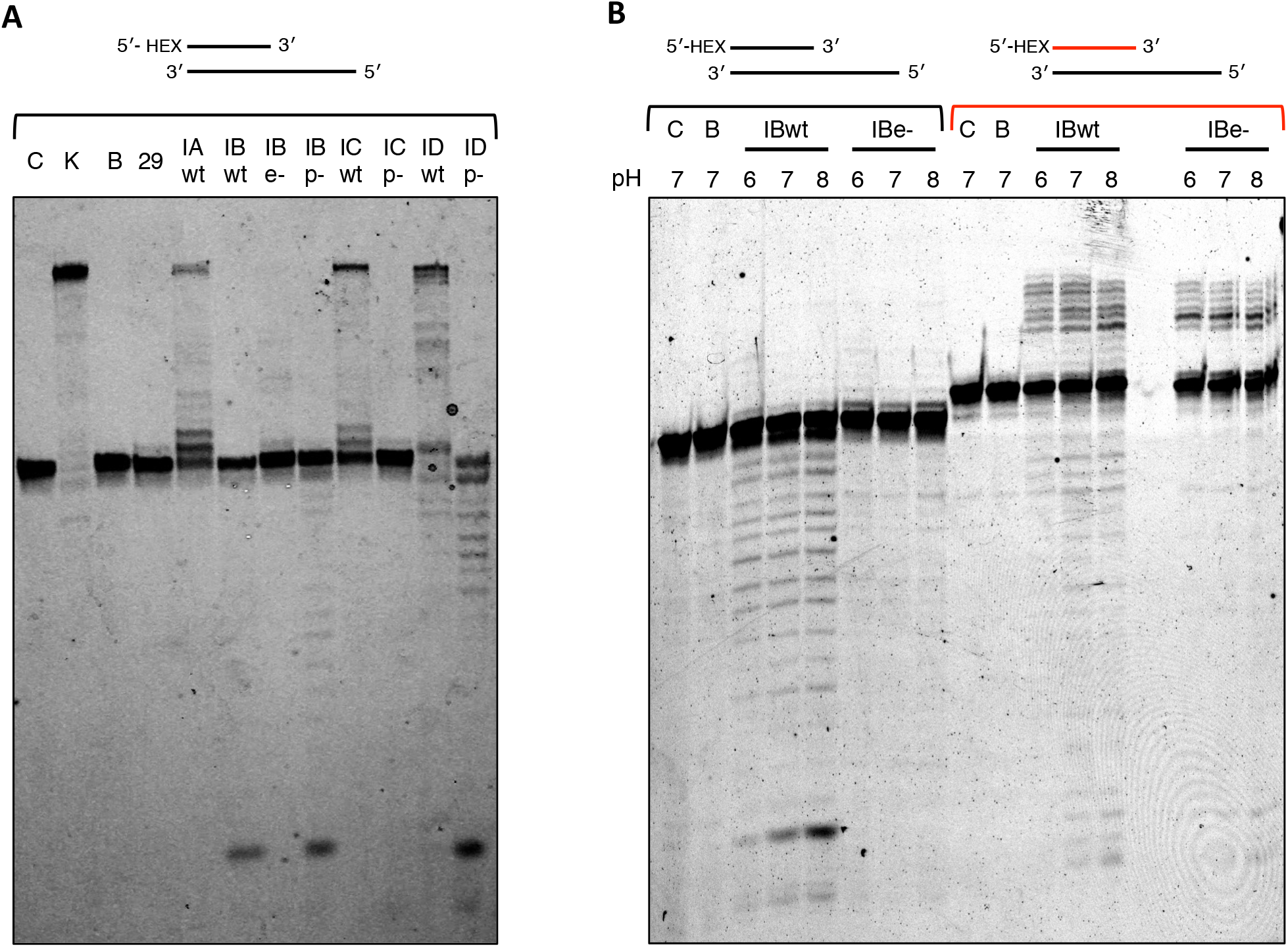
Pol and Exo activity of immunoprecipitated *T. brucei* Pol I-like paralogs. Gel-based extension assays were used to demonstrate activity of immunoprecipitated protein. (A) Each reaction contained 840 nM DNA primed substrate, 5 μL IgG beads bound to immunoprecipitated protein, using standard reaction buffer at pH 8.0 with 500 μM dNTPs for 60 minutes. No protein control Time=0, C; 1U Klenow fragment positive control, K; IgG beads control, B; 29-13 parental cells with IgG beads, 29. (B) POLIB variants in pol reactions at pH 6.0, 7.0, or 8.0, including 500 μM dNTPs with 840 nM either DNA primed template or RNA primed template for 60 minutes. Red oligo: RNA, Black oligo: DNA.

## DISCUSSION

The predicted structure of POLIB models is highly divergent from structures of all other characterized Family A Pols. In other well-characterized proteins in this family including *E. coli* Pol I, T7 DNA Pol, and Mammalian Pol g, have the exonuclease domain located underneath the pol domain. These *de novo* models of POLIB predict the 3’-5’ exonuclease domain to be inserted within the thumb of the pol domain (Figure 1, Figure S1). We have confidence in this arrangement of catalytic domains due to the generation of similar predictions in both RoseTTAfold and Alphafold (Figure 1B, S3) Based on the robust exonuclease activity of this enzyme that outcompetes pol activity under physiologically relevant conditions, it is possible the insertion interferes with the ability of the pol domain to carry out its DNA polymerase activity. This structural feature of POLIB was likely developed after the gene duplication event that gave rise to four Pol I-like proteins, giving rise to the specialized function of POLIB in kDNA replication.^6^

When we investigated the activity of the POLIB exo domain using recombinant protein, IBPol- consistently showed reduced exonuclease activity compared to wild-type protein (Figure 2). This suggests that there is crosstalk between the two domains, where a structural change in one impact the activity of the other. This is a phenomenon described in other pols such as human Pol g and may be a more dramatic effect in POLIB due to the divergent arrangement of the two functional domains.^28^

In recombinant IBWT, exo activity unexpectedly prevails over pol activity at higher pH conditions (Figure 5). These more basic conditions are likely closer to the physiological pH of the mitochondrial matrix, based on what has been reported in other systems. For example, in *Saccharomyces cerevisiae,* the cytosolic pH is ~7.2 under typical growth conditions while the mitochondrial matrix pH is more basic, ~7.5.^37^ Similarly, in HeLa cells, the pH of the cytosol is ~7.4 and the mitochondrial matrix pH is ~8.0.^38,39^ While the pH of the mitochondrial matrix of trypanosomes has not been experimentally determined, the pH of the cytosol has been reported to be 7.47± 0.06.^40^ We expect that the environment of the mitochondrial matrix in trypanosomes follows a similar pattern seen in the other organisms and is more basic than the cytosol. This suggests that POLIB exo activity on DNA substrates is dominant over pol activity in the alkaline *in vivo* environment.

Interestingly, Gluenz and colleagues defined discrete regions around the kDNA disk via electron microscopy and ethanolic phosphotungstic acid staining where basic proteins accumulated during kDNA replication.^41^ These regions likely correspond with the previously described antipodal sites and kinetoflagellar zones defined by immunofluorescence microscopy.^42,43^ If POLIB is recruited to these sites, it would likely be an optimal climate for exo activity based on the differential regulation of POLIB activities we demonstrated (Figure 5). This localization to particular environmental conditions could be a mechanism of regulating the balance between exo and pol activities for POLIB.

Trypanosomatids also contain two kDNA Pol β-like repair paralogs that differ in their kDNA associated localizations and have distinct optimal conditions, suggesting that they are also specialized for the conditions to which they localize.^4^ Additionally, the Pol I-like paralogs POLIC and POLID display spatiotemporal localization to the antipodal sites only during kDNA replication stages of the cell cycle representing another example of functional regulation of kDNA proteins via localization.^44,45^ POLIB also localizes near the kDNA, however a detailed study including the precise kDNA-associated region, and a cell cycle localization pattern has not yet been established.^5^ Similar to POLIC and POLID, POLIB may spatiotemporally localize to areas around the disc that have ideal conditions for one of the enzymatic activities or the other.

Another mechanism that could be altering POLIB activity is the use of different available divalent cations. Regulation by divalent cations has been described for other DNA Pols, such as human mitochondrial Pol γ, that has an enhanced ability to perform translesion synthesis (TLS) in the presence of Mn^2+^ compared to Mg^2+^.^46^ In another example, Pol η shifts its activity to ribonucleotide incorporation through an increased affinity to rNTPs in the presence of Mn^2+^.^47^ This modulation in activity for both proteins is likely due to a conformational change of the active site that is allowed due to the more relaxed coordination requirement of Mn^2+^.^48^ We show that POLIB extends more rapidly and with less fidelity when using Mn^2+^ compared to Mg^2+’^ suggesting the active site of POLIB may also be more accommodating to structural rearrangements in the presence of Mn^2+^ (Figure S11). For both Pol γ and Pol η, even a relatively low concentration of Mn^2+^ compared to Mg^2+^ in a reaction impacted the activity of the enzyme. Although the concentrations of Mg^2+^ and Mn^2+^ in the *T. brucei* mitochondrion have not been experimentally determined, both Mg^2+^ and Mn^2+^ are canonically found in mitochondria of other systems.^49,50,51,52^ We therefore expect that Mn^2+^ is present in the trypanosome mitochondrion, and that even in relatively small amounts, could impact POLIB *in vivo* to more rapidly extend DNA.

It is possible that POLIB activity is modulated *in vivo* by an accessory subunit, protein-protein interaction, or post-translational modification to improve pol activity. This kind of regulation could be mediated by two arginine methylation sites that have been identified in the N-terminal UCR domain of POLIB (Figure S1).^53^ These sites are not present in the truncated recombinant protein purified for the assays in this study. However, it is possible that the presence and methylation status of these residues play a part in regulating the activity of POLIB, either directly through some interdomain interaction on the protein or indirectly through facilitating protein-protein interactions or localization.

This work provides the first demonstration of enzymatic activity of all four mitochondrial Pol I-like proteins of *T. brucei*. One model to explain the presence of four A-Family pols in the mitochondrion of this system postulates that one protein acts as a replicative polymerase while the other three are playing support roles in kDNA maintenance. However, POLIB, POLIC, and POLID have been shown to be essential for kDNA replication by RNAi.^5,8^ In our assay of immunoprecipitated proteins, reactions containing POLIA, POLIC, and POLID resulted in primers that were extended to the end of the DNA template in higher proportions than the reaction products of POLIB (Fig 7A). Based on these data, the other three Pol I-like proteins are all more likely than POLIB to replicate DNA with high processivity, a defining feature of a replicative polymerase. This suggests that out of these polymerases, POLIB is not likely to be a replicative enzyme.

Another model to account for the three essential Family A pols proposes that the enzymes are working cooperatively similar to nuclear DNA replication in other model systems. Pol a from yeast demonstrates increased nucleotidyl incorporation activity from an RNA primer in comparison to a DNA primer, likely due to a higher affinity to A-form helices typically formed by RNA/DNA duplexes.^54^ In nuclear DNA replication, Pol a extends the RNA primers for completion of an RNA/DNA primer. Because POLIB extension activity is increased from an RNA primer (Figures 6,7) we hypothesize that it may be playing a similar role in trypanosomatid mitochondrial DNA replication to facilitate extension by a replicative pol.

When POLIB RNAi is induced, more linearized DNA minicircles are detected in Southern blots than in uninduced cells.^9^ Based on the robust exonuclease activity of POLIB that we have demonstrated herein, one explanation for this observation is that POLIB is degrading these linearized molecules *in vivo.* Data from other systems support this hypothesis. In human mitochondria, the exonuclease activity of Pol γ participates along with other exonucleases in the degradation of linearized DNA molecules.^55^ After the gene duplication event resulting in the four mitochondrial Pol I-like enzymes in *T. brucei,* POLIB may have retained this function of “cleaning up” molecules resulting from double-strand breaks. The preference for POLIB to degrade ssDNA may indicate that it is working in tandem with another enzyme that is degrading linearized molecules in the 5’-3’ direction, analogous to the function of MGME1 in humans.^55^

Because of the structural divergence of POLIB and its conservation across trypanosomatid organisms, POLIB is an attractive potential drug target for the treatment of diseases caused by medically relevant trypanosomatids. More studies are needed to address the essential role of POLIB *in vivo.* For example, complementation experiments can be done with POLIB enzymatic mutants to determine if the exonuclease activity is the essential contribution of POLIB to kDNA replication.

This study represents a new framework for the catalytic activity of a divergent Family A DNA polymerase, adding to the body of knowledge of how these diverse and important proteins can be functionally specialized. Our work also provides new insight into possible mechanisms of the complex process of kDNA replication in trypanosomatid organisms.

### ACCESSION CODES (Uniprot)

POLIA (Trypanosoma brucei) Q8MWB5

POLIB (Trypanosoma brucei) Q8MWB4

POLIC (Trypanossoma brucei) Q8MWB3

POLID (Trypanosoma brucei) Q8MWB2

POLIA (Arabidopsis thaliana) F4I6M1

POLIB (Arabidopsis thaliana) Q84ND9

TREX1 (Homo sapiens) Q9NSU2

TREX2 (Homo sapiens) Q9BQ50

5 (DNA- directed DNA polymerase) (Bacteriophage T7) P00581

PF3D7_1411400 (P. falciparum) Q8ILY1

PolA (E. coli) P00582

POLG (Homo sapiens) P54098

POLN (Homo sapiens) Q7Z5Q5

POLQ (Homo sapiens) O75417

POLE (Homo Sapiens) Q07864

BSAL_33270 (Bodo saltans) A0A0S4JJF3

### ACCESSION CODES for POLIB sequences (VEuPathDB Identifier Numbers)

CFAC1_220029600 (Crithidia fasciculata, Cf-cl)

EMOLV88_130005700 (Endotrypanum monterogeii)

LmjF.13.0080 (Leishmania major, Friedlin strain)

LtaP13.0080 (Leishmania tarentolae, Parrot-TarII strain)

Lsey_0383_0090 (Leptomonas seymouri)

PCON_0045170 (Paratrypanosoma confusum)

TcIL3000.11.4800 (Trypanosoma congolense)

TcCLB.506227.80 (Trypanosoma cruzi, CL Brener Esmeraldo-like strain)

TevSTIB805.11_01.4840 (Trypanosoma evansi)

## Supporting information

All supplemental data

## ASSOCIATED CONTENT

### Supporting Information

Scripts used for Dynafit global fitting of processivity progress curves, alignments of amino acid sequences of POLIB and other Family A DNA polymerases, Alphafold structure of POLIB and data on quality of predicted protein structures, nucleotide sequence of codon-optimized recombinant POLIB, oligos used for cloning and site directed mutagenesis, table of T*rypanosoma brucei* cell lines used for this study, additional experimental data including representative purification of recombinant POLIB, enzymatic assays, and results of immunoprecipitations are all included in a supplementary PDF file.

## AUTHOR INFORMATION

### Author Contributions

M.M.K., S.B.D., and S.W.N. designed experiments. S.B.D. and M.P.F. performed experiments. M.M.K., S.W.N. and S.B.D. discussed results. The manuscript was written through contributions from all authors. All authors have given approval to the final version of the manuscript.

### Notes

The authors declare no competing financial interests.

## ACKNOWLEDGMENT

The authors thank Drs. Peter Chien, Klaus Nüsslein, and Derek Lovley for generously sharing equipment; Dr. Jeanne Hardy for help in protein purification protocol design; and members of our research group for their valuable comments and feedback throughout the conduct of this study and preparation of the manuscript. We would also like to thank Raveen Armstrong, Dr. Minu Chaudhuri, Dr. Craig Martin, and Dr. Jonathan Miller for helpful comments on the manuscript. Access to the ÄKTA pure chromatography system and Typhoon™ FLA 9500 imager (GE) was through the Core facilities in the Institute for Applied Life Sciences (University of Massachusetts Amherst).

## ABBREVIATIONS

kDNA: kinetoplast DNA;
Pol: polymerase;
Exo: exonuclease;
dNTPs: deoxyribonucleotide triphosphates;
rNTPs: ribonucleotide triphosphates;
DTT: dithiothreitol;
EDTA: ethylenediaminetetraacetic acid;
HEX: hexachlorofluorescein,
SD: Standard deviation;
BSA: bovine serum albumin,
TLS: translesion DNA synthesis,
IBWT: TbPOLIB truncated recombinant protein;
IBPol-: TbPOLIB recombinant protein with polymerase catalytic active site mutations;
IBExo-: TbPOLIB recombinant protein with exonuclease catalytic active site mutations.

