## Supplementary material for "Trypanosoma brucei Mitochondrial DNA Polymerase POLIB Contains a Novel Thumb Insertion That Confers Dominant Exonuclease Activity": All supplemental data

ABSTRACT

Trypanosoma brucei and related parasites contain an unusual catenated mitochondrial genome known as kinetoplast DNA (kDNA) composed of maxicircles and minicircles. The kDNA structure and replication mechanism are divergent and essential for parasite survival. POLIB is one of three Family A DNA polymerases independently essential to maintain the kDNA network. However, the division of labor among the paralogs, particularly which might fulfill the role of a replicative, proofreading enzyme remains enigmatic. De novo modeling of POLIB revealed a structure divergent from all other Family A polymerases in which the thumb subdomain contains a 369 amino acid insertion with homology to DEDDh DnaQ family 3'-5' exonucleases. Here we demonstrate recombinant POLIB 3'-5' exonuclease prefers DNA vs. RNA substrates and degrades single- and double-stranded DNA in a non-processive manner. The exonuclease activity prevails over polymerase activity on DNA substrates at pH 8.0, while DNA primer extension is favored at pH 6.0. Mutations that ablate POLIB polymerase activity slow the exonuclease rate suggesting crosstalk between the domains. We show that POLIB is able to extend an RNA primer more efficiently than a DNA primer in the presence of dNTPs but does not incorporate rNTPs efficiently using either primer. Immunoprecipitation of Pol I-like paralogs from T. brucei corroborate the pH selectivity and RNA primer preferences of POLIB and revealed that the other paralogs efficiently extend a DNA primer. The unique POLIB thumb insertion influences the balance between polymerase and exonuclease activity and provides another example of exquisite diversity among DNA polymerases for specialized function.

```
[task]
task = fit
data = progress
model = 2step?
[mechanism]
E*DDDDDDDDDDDDDDDDDDDDDD <===> E*  +  DDDDDDDDDDDDDDDDDDDDDD  : k-2 k1
E*DDDDDDDDDDDDDDDDDDDDDD --> EDDDDDDDDDDDDDDDDDDDDDD : kconf
EDDDDDDDDDDDDDDDDDDDDDD --> EDDDDDDDDDDDDDDDDDDDDD + R   : kexo
EDDDDDDDDDDDDDDDDDDDDD <==> E  +  DDDDDDDDDDDDDDDDDDDDD  : k-2 k1
EDDDDDDDDDDDDDDDDDDDDD --> EDDDDDDDDDDDDDDDDDDDD + P   : kexo
EDDDDDDDDDDDDDDDDDDDD <==> E  +  DDDDDDDDDDDDDDDDDDDD  : k-2 k1
EDDDDDDDDDDDDDDDDDDDD --> EDDDDDDDDDDDDDDDDDDD + P   : kexo
EDDDDDDDDDDDDDDDDDDD <==> E  +  DDDDDDDDDDDDDDDDDDD  : k-2 k1
EDDDDDDDDDDDDDDDDDDD --> EDDDDDDDDDDDDDDDDDD + P   : kexo
EDDDDDDDDDDDDDDDDDD <==> E  +  DDDDDDDDDDDDDDDDDD  : k-2 k1
EDDDDDDDDDDDDDDDDDD --> EDDDDDDDDDDDDDDDDD + P   : kexo
EDDDDDDDDDDDDDDDDD <==> E  +  DDDDDDDDDDDDDDDDD  : k-2 k1
EDDDDDDDDDDDDDDDDD --> EDDDDDDDDDDDDDDDD + P   : kexo
EDDDDDDDDDDDDDDDD <==> E  +  DDDDDDDDDDDDDDDD  : k-2 k1
EDDDDDDDDDDDDDDDD  --> EDDDDDDDDDDDDDDD + P   : kexo
EDDDDDDDDDDDDDDD <==> E  +  DDDDDDDDDDDDDDD  : k-2 k1
EDDDDDDDDDDDDDDD --> EDDDDDDDDDDDDDD + P   : kexo
EDDDDDDDDDDDDDD <==> E  +  DDDDDDDDDDDDDD  : k-2 k1
EDDDDDDDDDDDDDD --> EDDDDDDDDDDDDD + P   : kexo
EDDDDDDDDDDDDD <==> E  +  DDDDDDDDDDDDD  : k-2 k1
EDDDDDDDDDDDDD --> EDDDDDDDDDDDD + Q   : kexo
EDDDDDDDDDDDD <==> E  +  DDDDDDDDDDDD  : k-2 k1
EDDDDDDDDDDDD --> EDDDDDDDDDDD + P   : kexo
EDDDDDDDDDDD <==> E  +  DDDDDDDDDDD  : k-2 k1
EDDDDDDDDDDD --> EDDDDDDDDDD + P   : kexo
EDDDDDDDDDD <==> E  +  DDDDDDDDDD  : k-2 k1
EDDDDDDDDDD  --> EDDDDDDDDD + P   : kexo
EDDDDDDDDD <==> E  +  DDDDDDDDD  : k-2 k1
EDDDDDDDDD  --> EDDDDDDDD + P   : kexo
EDDDDDDDD <==> E  +  DDDDDDDD  : k-2 k1
EDDDDDDDD  --> EDDDDDDD + P   : kexo
EDDDDDDD <==> E  +  DDDDDDD  : k-2 k1
EDDDDDDD  --> EDDDDDD + P   : kexo
EDDDDDD <==> E  +  DDDDDD  : k-2 k1
EDDDDDD  --> EDDDDD + P   : kexo
EDDDDD <==> E  +  DDDDD  : k-2 k1
EDDDDD  --> EDDDD + P   : kexo
EDDDD <==> E  +  DDDD  : k-2 k1
EDDDD  --> EDDD + P   : kexo
EDDD <==> E  +  DDD : k-2 k1
EDDD  --> EDD + P   : kexo
EDD <==> E  +  DD : k-2 k1
EDD  --> E  +  D  +  P   : kexo
[constants]
k1 = 50
kexo = 1??
k-2 = 34??
kconf = 1??
```

**Figure S1. Dynafit script for global fitting of 1AP and 2AP progress curves for the ssDNA and blunt dsDNA substrates.**

```
[task]
task = fit
data = progress
model = 2step?
[mechanism]
EDDDDDDDDDDDDDDDDDDDDDD <===> E  +  DDDDDDDDDDDDDDDDDDDDDD  : k-2 k1
EDDDDDDDDDDDDDDDDDDDDDD -->  EDDDDDDDDDDDDDDDDDDDDD + R   : kexo
EDDDDDDDDDDDDDDDDDDDDD <==> E  +  DDDDDDDDDDDDDDDDDDDDD  : k-2 k1
EDDDDDDDDDDDDDDDDDDDDD -->  EDDDDDDDDDDDDDDDDDDDD + P   : kexo
EDDDDDDDDDDDDDDDDDDDD <==> E  +  DDDDDDDDDDDDDDDDDDDD  : k-2 k1
EDDDDDDDDDDDDDDDDDDDD -->  EDDDDDDDDDDDDDDDDDDD + P   : kexo
EDDDDDDDDDDDDDDDDDDD <==> E  +  DDDDDDDDDDDDDDDDDDD  : k-2 k1
EDDDDDDDDDDDDDDDDDDD -->  EDDDDDDDDDDDDDDDDDD + P   : kexo
EDDDDDDDDDDDDDDDDDD <==> E  +  DDDDDDDDDDDDDDDDDD  : k-2 k1
EDDDDDDDDDDDDDDDDDD -->  EDDDDDDDDDDDDDDDDD + P   : kexo
EDDDDDDDDDDDDDDDDD <==> E  +  DDDDDDDDDDDDDDDDD  : k-2 k1
EDDDDDDDDDDDDDDDDD -->  EDDDDDDDDDDDDDDDD + P   : kexo
EDDDDDDDDDDDDDDDD <==> E  +  DDDDDDDDDDDDDDDD  : k-2 k1
EDDDDDDDDDDDDDDDD -->  EDDDDDDDDDDDDDDD + P   : kexo
EDDDDDDDDDDDDDDD <==> E  +  DDDDDDDDDDDDDDD  : k-2 k1
EDDDDDDDDDDDDDDD -->  EDDDDDDDDDDDDDD + P   : kexo
EDDDDDDDDDDDDDD <==> E  +  DDDDDDDDDDDDDD  : k-2 k1
EDDDDDDDDDDDDDD -->  EDDDDDDDDDDDDD + P   : kexo
EDDDDDDDDDDDDD <==> E  +  DDDDDDDDDDDDD  : k-2 k1
EDDDDDDDDDDDDD -->  EDDDDDDDDDDDD + Q   : kexo
EDDDDDDDDDDDD <==> E  +  DDDDDDDDDDDD  : k-2 k1
EDDDDDDDDDDDD -->  EDDDDDDDDDDD + P   : kexo
EDDDDDDDDDDD <==> E  +  DDDDDDDDDDD  : k-2 k1
EDDDDDDDDDDD -->  EDDDDDDDDDD + P   : kexo
EDDDDDDDDDD <==> E  +  DDDDDDDDDD  : k-2 k1
EDDDDDDDDDD -->  EDDDDDDDDD + P   : kexo
EDDDDDDDDD <==> E  +  DDDDDDDDD  : k-2 k1
EDDDDDDDDD  -->  EDDDDDDDD + P   : kexo
EDDDDDDDD <==> E  +  DDDDDDDD  : k-2 k1
EDDDDDDDD  -->  EDDDDDDD + P   : kexo
EDDDDDDD <==> E  +  DDDDDDD  : k-2 k1
EDDDDDDD  -->  EDDDDDD + P   : kexo
EDDDDDD <==> E  +  DDDDDD  : k-2 k1
EDDDDDD  -->  EDDDDD + P   : kexo
EDDDDD <==> E  +  DDDDD  : k-2 k1
EDDDDD  -->  EDDDD + P   : kexo
EDDDD <==> E  +  DDDD  : k-2 k1
EDDDD  -->  EDDD + P   : kexo
EDDD <==> E  +  DDD : k-2 k1
EDDD  -->  EDD + P  : kexo
EDD <==> E  +  DD : k-2 k1
EDD  -->  E + D + P : kexo
E + P <==>  EP : k1 k-3;
[constants]
k1 = 50
kexo = 1??
k-2 = 34??
k-3 =5??
```

**Figure S2. Dynafit script for global fitting of 1AP and 2AP progress curves for the recessed dsDNA substrate.**

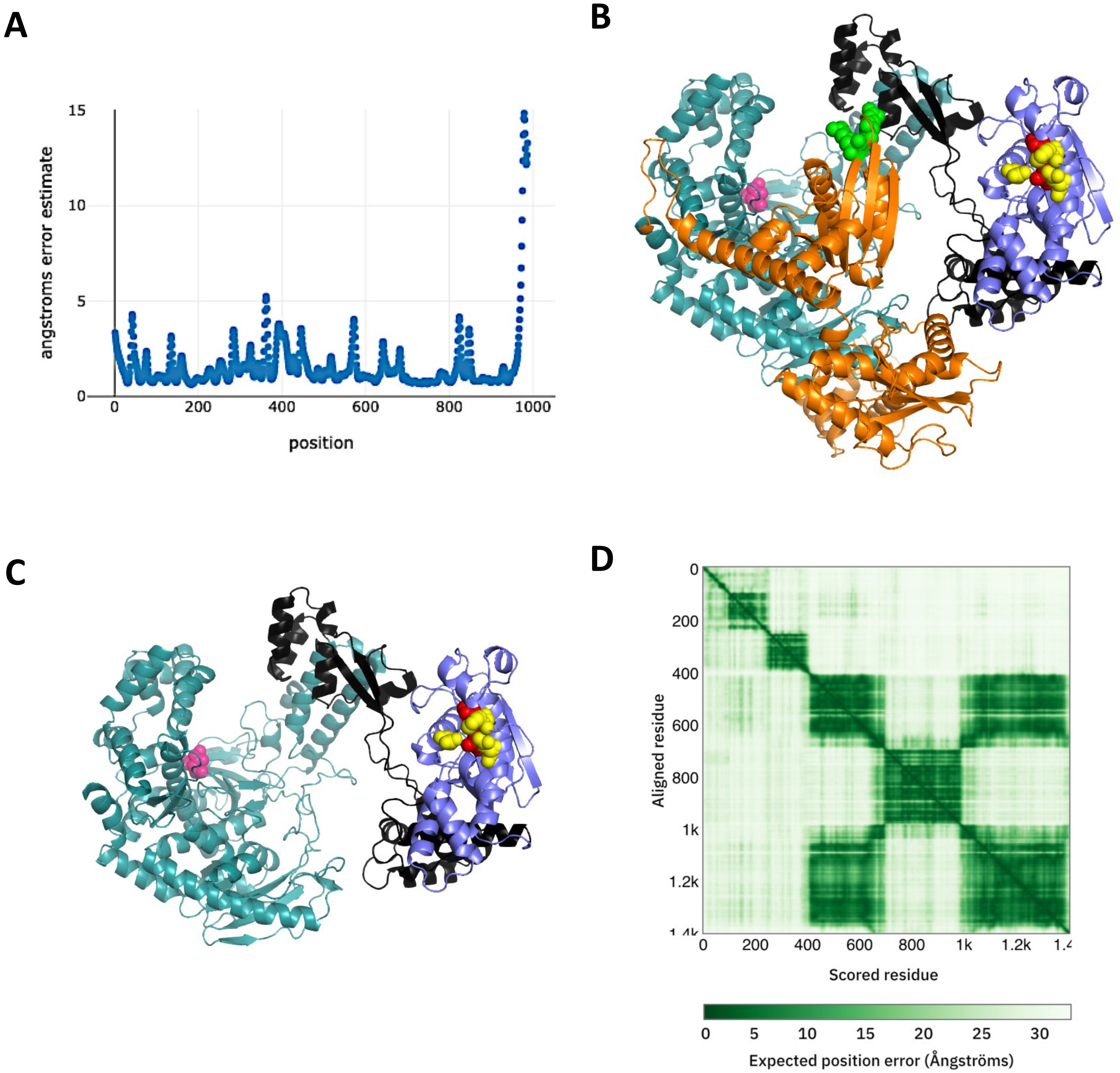


**Figure S3. Quality of Rosettafold model and comparison of domain structure with Alphafold (A)** Per Residue Error Estimate of TbPOLIB RoseTTAfold Model displayed in Figure 1. **(B)** Alphafold Model of Full-length POLIB. Model retrieved from Alphafold database (Entry: Mitochondrial DNA pol I protein B, Tb11.02.2300, *Trypanosoma brucei brucei* (strain 927/4 GUTat10.1).[56] Teal, pol domain; purple, portion of insertion with homology to an exonuclease domain; black, thumb insertion linker regions; Magenta, pol domain active sites mutated for IBpol- variant, red, exo catalytic residues for IBexo- variant; yellow, additional exonuclease residues of the DEDDh motif. **(C)** Alphafold model of POLIB (residues 414-1400). Same color scheme as in B. Orange, uncharacterized C-terminal domain (UCR); bright green, arginine methylation sites (R266, R270).[53] **(D)** Predicted aligned error for full length Alphafold model.

```
E. coli Pol I                ----------    -------------------------------    --------------------------------
H. sapiens Pol γ     650     RAIESLYRKH    ----CLEQGKQQLMPQEAGLAEEF---LLTD    -NSAIWQTVEELDYLEVEAEAKMENLRAAVp   713
T7 DNA Pol           8       ESLEAVDIEH    RAAWLLAKQERNGFPFDTKAIEELYVELAAR    -RSELLRKLTETFGSWYQPKGGTEMFCHPRt   78
H. sapiens Pol θ     475     HELPLLEGME    ------TSQGIQSLGLNAGSEHSGRYRASVE    -SILIFNSMNQLNSLLQK-----ENLQ----   530
H. sapiens Pol ν     368     -----LVEKY    --------CEK-SITVKVNSTYGNSSRNIV-    -NQNVRENLKTLYRLTMDLCSKLKDYGL---   420
T. brucei POLIA      399     -----LLQSY    --------QQQIGLPAGLNAAECKGSKEAV-    -SHRVFY-MEPLYRM---LYGRLGSHGL---   448
T. brucei POLIB      710     HLCHKYGN--[ 4]SADVHLQR----------FASERGLRGAGR[14]SRRRKYRLVVFDIESTGLNTATDAIIEVAA-   785
T. brucei POLIC      926     HLGTRYVSPH[16]LTTTPVPE-------LYTNAENLLGVKGTGN[36]LDNLLNNTAIVVVTCKVRTSSFLEVITIHC-   1041
T. brucei POLID      817     ERHTLYARPE[27]ATGVSLDKGTKFFKMYFEAYVKQAGFRKSSS[48]ISKLFLKNAITGESLTIAGDCANKVERLRA-   962
A. thaliana PolIB    418     PFGELLVKME    AEGILVDREYLAEIEKVAKAEQQVAGSRFRN[ 5]CPDAKYMNIGSDTQLRQLFFGGISNS-H---   490
A. thaliana PolIA    494     PFGELLAKME    SEGMLVDRDYLAQIEIVAKAEQEIAVSRFRN[ 5]CPDAKHMNVGSDTQLRQLFFGGISNSCN---   567
T. gondii Prex       1833    ------SSSE[20]SGGAAGDDDGSDEREVRLSMGVDVETTGLDP[43]IRKVFHNGQFDLCFLAAAGLADRDAKRQEVf   1961

E. coli Pol I        1       -GPLNVFENIEMPLVPV    LSRIERNGV-KIDPKVLHNHSEELTLRLAELEKKAHEI--A    GEEFNLSSTKQLQT   68
H. sapiens Pol γ     714     GQPL------------A    LTARGGPKDTQPSYHHGNGPYNDVDIPGC-WFFKLPHK--D    GNSCNVGSPFAKDF   770
T7 DNA Pol           79      GKPLPKYPRIKTPKVGG    IFKKPKNKAQREGREPCELDTREYVA-----GAPYTPV--E    HVVFNPSSRDHIQK   143
H. sapiens Pol θ     531     ----DVFRKVEMPSQYC    LALLELNGI-GFSTAECESQKHIMQAKLDAIETQAYQL--A    GHSFSFTSSDDIAE   595
H. sapiens Pol ν     421     ---WQLFRTLELPLIPI    LAVMESHAI-QVNKEEMEKTSALLGARLKELEQEAHFV--A    GERFLITSNNQLRE   486
T. brucei POLIA      449     ---LPAFLKQEKRISIM    CAAMKLNGF-YVNLVEVDGFKARCAEKMEKLRSEARSMipS    MTNFNIQSADDCRV   516
T. brucei POLIB      786     FDPVEGTSFSSLV----[52]LIGHNVFALDEPLLRRAFRSESVDTENLLFCDSLTIL----[ 1]GLKQEL--QGSKKD   900
T. brucei POLIC      1042    PSSDPNDELRHTVAIDD[58]LVDTGIIQTIAETVLGKENEERSLLERVCFCDINRAFPpp-[12]GLVNGAEQAGVSSR   1182
T. brucei POLID      963     FMEGQVEAHTEMLVDAD[60]TLADTVSTLSYYRYMYGSDTEAAYADKTKVDESKLVVPmkA[13]AFKKKLVQEAAMKP   1107
A. thaliana PolIB    491     DEVLPVEKLFKVPNIDK    VIEEGKKTPTKFRNIKLHRISDSPLSTENFTASGWPSV--S    GDVLKELAGKVSAE   560
A. thaliana PolIA    568     DEDLPYEKLFKVPNVDK    VIEEGKKRATKFRNIKLHRISDRPLPTEKFTASGWPSV--S    GDTLKALAGKVSAE   637
T. gondii Prex       1962    SESQDAPESQSRPHFPT[64]LLDAAVLGARPTEVGDLQSDGGVFVSGPLFDTLIAAKVveA[13]ERFLGVLMDKRMQA   2110

E. coli Pol I        69      ILFEKQGIKP---------------LKKTPGGAP----    -----------STSEEVLEELALD--YPLPKVILEY      110
H. sapiens Pol γ     771     LPKMEDGTLQ---------------AGPGGASGPR---    -----------ALEINKMISFWRN-AHKRISSQMVV[1]   815
T7 DNA Pol           144     KLQEAGWV-P---------------TKYTDKGAP----    -----------VVDDEVLEGVRVDDPEKQAAIDLIK[4]   190
H. sapiens Pol θ     596     VLFLELKLPPNREMK-------NQGSKKTLGSTRRGID    NGRKLRLGrqfSTSKDVLNKLKAL--HPLPGLILEW      660
H. sapiens Pol ν     487     ILFGKLKL--HLLS-------QRNSLPRTGLQKYP---    -----------STSEAVLNALRDL--HPLPKIILEY      535
T. brucei POLIA      517     ALYEVLKLGPNLVSKGTEGSTENNSLILTRGGK-L---    -----------STSEETLRILARH--HEFPRILIAY      573
T. brucei POLIB      901     SKFDRGVLDILTNSLRLSSLVEGLRVEADGELHRADTD[ 66]EKRTIEQLrkrHLDEATFVVLQR-HKLEVAGLLLQK   1039
T. brucei POLIC      1183    SNW-EAFTRNFAAVGGFEDTRRALVLHCASLVGTARDP[ 62]ERINVLLSnekQVDLSFFKALQQ---LRFMEKQMQL   1314
T. brucei POLID      1108    EDWCNLYIDIFLHIGHAALCRESLETEAGSKKARKSK-[ 85]RKFRSPTTrqlQVGEDSLTYFKKTHGDKTAETILEL   1265
A. thaliana PolIB    561     YDFTDDVSDISLEEVVEDDDV-----ETSETQKSKTDD    ETDTSAYG---TAYVAFGGGERGKEACHAIASLCEV      626
A. thaliana PolIA    638     YDYMEGVLDTCLEENIGDDDCISLPDEVVETQHVNTSV    ESDTSAYG---TAFDAFGGGESGKEACHAIAALCEV      708
T. gondii Prex       2111    SDWSSPHLSQEQLLYAARDAAVLLPLQQRLQQKLEAFD[101]FDKNAAEVedqLLKDTSDGTLARLSQFPAVQALRDY   2285

E. coli Pol I        111     RGLAKLKSTYTDKLPLM    I--NPKTG    RVHTSYHQAVTATGRLSSTDPNLQNIP---[ 1]RNEEGR    RIRQ   171
H. sapiens Pol γ     816     LPRSALPRAVIRHPDYD    E--EGLYG    AILPQVVTAGTITRRAVEPTWLTASNA---[ 1]PDRVGS    ELKA   876
T7 DNA Pol           191     IQKRIGQSAEGDKAWLR    Y--VAEDG    KIHGSVNPNGAVTGRATHAFPNLAQIPG--[ 1]RSPYGE    QCRA   252
H. sapiens Pol θ     661     RRITNAITKVVFPLQRE    KCLNPFLG[ 2]RIYP-VSQSHTATGRITFTEPNIQNVPR--[ 6]PTLVGE[46]SMRH   776
H. sapiens Pol ν     536     RQVHKIKSTFVDGLL--    --ACMKKG    SISSTWNQTGTVTGRLSAKHPNIQGISKHP[ 7]-NFKGK[ 7]SPRA   609
T. brucei POLIA      574     RKLAKLLQTYVVGFMEM[1]IPIEEGDG[27]KLHPNLVQEGTETGRLTSVEPNMQALPRST[14]DSENGN[28]FIRR   708
T. brucei POLIB      1040    LQLERGSANFLHSGTDG[1]LSILHSDN    KVRQYIDLTATTTSRTTSSYPSCQNIPKDD    ------[ 2]SLRH   1101
T. brucei POLIC      1315    FEEGALFRAVLP-----    ----ECKN    RVHGEFCHVVTATGRLSSQSPNLQNIPKED    ------    -LRC   1363
T. brucei POLID      1266    RAMEKLIGTYYENTDDG[4]SLVHDDDS    CIHHELIHNKTNTGRLASANPNCQNIPKED    ------[ 2]PLRE   1330
A. thaliana PolIB    627     CSIDSLISNFILPLQGS    -NVSGKDG    RVHCSLNIN-TETGRLSARRPNLQNQPALE    KDRY--    KIRK   687
A. thaliana PolIA    709     CSIDSLISNFILPLQGS    -NVSGKDG    RVHCSLNIN-TETGRLSARRPNLQNQPALE    KDRY--    KIRQ   769
T. gondii Prex       2286    RKAAKAITTFVDKLPEH    --INSVTG    KIHCSLHQCGAGSGRFSCDSPNLQQIPRER    ------    RFRA   2342
```

Motif A

```
E. coli Pol I         172   AFIAPE    D-YVIVSADYSQIELRIMAHLSRDKGLLTAFAEGK--DIHRA     TAAEVFGLPLET    VT        SEQRR   235
H. sapiens Pol γ      877   MVQAPP    G-YTLVGADVDSQELWIAAVLGDAHFAGMHGCTAF--GWMTL     QGRKSRGTDLHS[4]TV        GISRE   944
T7 DNA Pol            253   AFGAEH[2]D-GITGKPWVQAGIDASGLELRCLAHFMARFDNGE--YAHEI     LNGDIHTKNQIA    AE        LPTRD   318
H. sapiens Pol θ      777   AFVPFP    G-GSILAADYSQLELRILAHLSHDRRLIQVLNTGA--DVFRS     IAAEWKMIEPES    VG        DDLRQ   840
H. sapiens Pol ν      610   MFVSSK    G-HTFLAADFSQIELRILTHLSGDPELLKLFQESErdDVFST     LTSQWKDVPVEQ    VT        HADRE   675
T. brucei POLIA       709   CLGVPD    G-YSLVSLDYEQVELRVLAHLSGDSALISVLTKSG--DIHRS     IAEIIFRKT--S    VT        GEERS   770
T. brucei POLIB      1102   LFVSRF[2]K-GRCVEIDYSQLEIVVMAVLCEDERLVSDLNQGV--DFHVK     RASFFSGISYDE    IY[10]LKLRK   1177
T. brucei POLIC      1364   LIVSRF[2]RaGRMIEADYSQLEVVVLAALSRDARMLQELNDNV--DFHCL     RVSLMTKEPYED    VI[11]IQLRQ   1441
T. brucei POLID      1331   MFVSRY[2]K-GMCIEADYSQLEVVALAVLANDQQMLEDLRSNV--DFHCK[9]QYTEILKKAKKD    K-[ 4]VKLRQ   1408
A. thaliana PolIB     688   AFVASP    G-NTLVVADYGQLELRILAHLTGCKSMMEAFKAGG--DFHSR     TAMNMYPHVREA    VE[26]GSERR   777
A. thaliana PolIA     770   AFIASP    G-NSLIVADYGQLELRILAHLASCESMKEAFIAGG--DFHSR     TAMNMYPHIREA    VE[26]ASERR   859
T. gondii Prex       2343   CFVPSK[5]P-GKFIIADFSQIELRIAADLACDERMIEAYRKGE--DLHRL     TASLILNKPPSL    LS        KADRQ  2411
```

Motif B

```
E. coli Pol I         236   SAKAINFGLIYGMSAFG    LARQLN    IPRKEAQKYMDLYFERYPGV    LEYMERTRA    QAKEQGYVETLDG       300
H. sapiens Pol γ      945   HAKIFNYGRIYGAGQPF    AERLLM[5]LTQQEAAEKAQQMYAATKGL[34]KVQRETARK[11]RAWKGGTESEMFN    1059
T7 DNA Pol            319   NAKTFIYGFLYGAGDEK    IGQIVG    AGKERGKELKKKFLENTPAI    AALRESIQQ[ 4]SSQWVAGEQQVKW   387
H. sapiens Pol θ      841   QAKQICYGIIYGMGAKS    LGEQMG    IKENDAACYIDSFKSRYTGI    NQFMTETVK    NCKRDGFVQTILG       905
H. sapiens Pol ν      676   QTKKVVYAVVYGAGKER    LAACLG    VPIQEAAQFLESFLQKYKKI    KDFARAAIA    QCHQTGCVVSIMG       740
T. brucei POLIA       771   LAKKVVFGILYGAGPRG    LAQQMG    VSVEQALRVSSLFKSCFPTV    DAYQRRIID    QCRSDGSVRTLSG       835
T. brucei POLIB      1178   VAKTFSFQRLYGAGVPL    LHKTTG    IPVQDLQECIRREEEEYPGI    SRFHRLART[ 5]NNPGLPTHFIVE-[ 9]  1255
T. brucei POLIC      1442   QAKTFSFQRQYGAGTST    IATTTG    LSETEVVRLIAAEEQHYKDL    GRYYRLVTD[ 4]GADRLLQLRTLDA[12]  1522
T. brucei POLID      1409   QAKIFSFQRQYGAGVKM    ISESTG    LTQDQVRHLIEKERETYRGV    DVFNSMVAL[11]GSRNVRGHQIFKG[12]  1496
A. thaliana PolIB     778   KAKMLNFSIAYGKTAVG    LSRDWK    VSTKEAQETVDLWYNDRQEV    RKWQEMRKK    EATEDGYVLTLLG       842
A. thaliana PolIA     860   KAKMLNFSIAYGKTAIG    LSRDWK    VSREEAQDTVNLWYNDRQEV    RKWQELRKK    EAIQKGYVLTLLG       924
T. gondii Prex       2412   LAKAVNFGLIYGMSADR[4]ASSAYG[2]MSLQEARDFHAKYFSSYPGI    TRWHRRQKA    --EQPRETRTRAG      2480
```

```
E. coli Pol I         301   RRL-     YLPDIKSS     NGARRAAAERA    AINAPMQGTAADIIKRAMIAVDAWLQAE---[ 3]------VRMIMQV   360
H. sapiens Pol γ     1060   KLES[4]DIPRTPVL[4]SRALEPSAVQE[5]RVNWVVQSSAVDYLHLMLVAMKWLFEEF---[ 3]G------RFCISI  1133
T7 DNA Pol            388   KRR-     WIKGLDGR     KVHVRSP--HA    ALNTLLQSAGALICKLWIIKTEEMLVEK---[ 3]HgWdgdFAYMAWV   451
H. sapiens Pol θ      906   RRR-     YLPGIKDN     NPYRKAHAERQ    AINTIVQGSAADIVKIATVNIQKQLETFHST[32]GgF-----FILQL   998
H. sapiens Pol ν      741   RRR-     PLPRIHAH     DQQLRAQAERQ    AVNFVVQGSAADLCKLAMIHVFTAVAASHTL        -----tARLVAQI   801
T. brucei POLIA       836   RVR-     SIPDINDR     VLTKRSHAERQ    AFNTVVQGSAADVMKLGMIAVEREVLQPHAP        -----dVRLLLQV   896
T. brucei POLIB      1256   KTRD     -VV-----     -----LNLPP-[1]-KNYPIQSFGAELAQMMIGRVFRQFVRKSFY        GqK---AFMINFV  1307
T. brucei POLIC      1523   LTEP[4]VVPTG---[3]DFTKDKKAVPR[1]-KNYPVQGLAGEIVQIMCGKIIRRFYAKRNY        NdK---AFLVNTV  1590
T. brucei POLID      1497   TESD     -VPEGLLR[3]LAVKSTNFSPT[2]-KNYPVQGFAGEIVQIMLGVLWRHFLRKDNY        GgL---AVLINTV  1563
A. thaliana PolIB     843   RSRR     -FPASKSR     --AQRNHIQRA    AINTPVQGSAADVAMCAMLEISTNQQLKKLG        --W----RLLLQI   900
A. thaliana PolIA     925   RARK     -FPEYRSR     --AQKNHIERA    AINTPVQGSAADVAMCAMLEISNNQRLKELG        --W----KLLLQV   982
T. gondii Prex       2481   ----     --------     ---RRALFEY[5]SLNYPIQGTSADITKESLVQLQHKLAPLGGR        --------LVMCV  2528
```

Motif C

```
E. coli Pol I         361   HDELVFEVHKDD--VDAVAKQIHQLME    NCTRL--     DVPLLVEVGSGENW-DQAH       ---          408
H. sapiens Pol γ     1134   HDEVRYLVREEDryRAALALQITNLLT    RCMFA--     YKLGLNDLPQSVAF-FSAV[24]RRY[29]     1239
T7 DNA Pol            452   HDEIQVGCRTEE-iAQVVIETAQEAMR    WVGDHwN[1]RCLLDTEGKMGPNW-AICH       ---          503
H. sapiens Pol θ      999   HDELLYEVAEED--VVQVAQIVKNEME    SAVKL--     SVKLKVKVKIGASW-GELK[ 4]---          1050
H. sapiens Pol ν      802   HDELLFEVEDPQ--IPECAALVRRTME[5]QALEL-Q[1]QVPLKVSLSAGRSW-GHLV[23]APG[18]     900
T. brucei POLIA       897   HDEIILSVPNHM--LHSIVPAAMHAF-    -AHPI-S[1]LVPLLVTTKVGRLL-GDL-[12]VPS          958
T. brucei POLIB      1308   HDSLWLDCHMSV--LEECVHETRTIME    EVDTYvA[8]KVPLKVSVDCGVDM-CAME[24]PEL[12]     1404
T. brucei POLIC      1591   HDCVWIDAHESV--ADEVMDDVSAIMS    STSEViS[8]DVPFKAEIHIGPSL-----       GEL[ 2]     1649
T. brucei POLID      1564   HDCVWIDCHMDV--LQDVVLETDGIMS    SVRDV--     LNRLYPEMEVSVDFpCDVV[ 9]PVV[ 5]     1629
A. thaliana PolIB     901   HDEVILEGPIES--AEIAKDIVVDCMS[4]GRNIL--     SVDLSVDAKCAQNW-YAAK       ---          952
A. thaliana PolIA     983   HDEVILEGPSES--AENAKDIVVNCMS[4]GKNIL--     SVDLSVDAKCAQNW-YAGK       ---         1034
T. gondii Prex       2529   HDEIIAEVPEEK--AEEGLRVLIDTME    AAGNKyL[1]FVPCVAEGAIADSW-ADKP       ---         2579
```

**Figure S4. Sequence alignment of Family A Polymerase Domain.**

COBALT Multiple sequence alignment of the POLA domain from selected Family A DNA polymerases. Essential catalytic residues are indicated in motifs I and III (red circles). Motifs A, B and C are indicated with a black line.

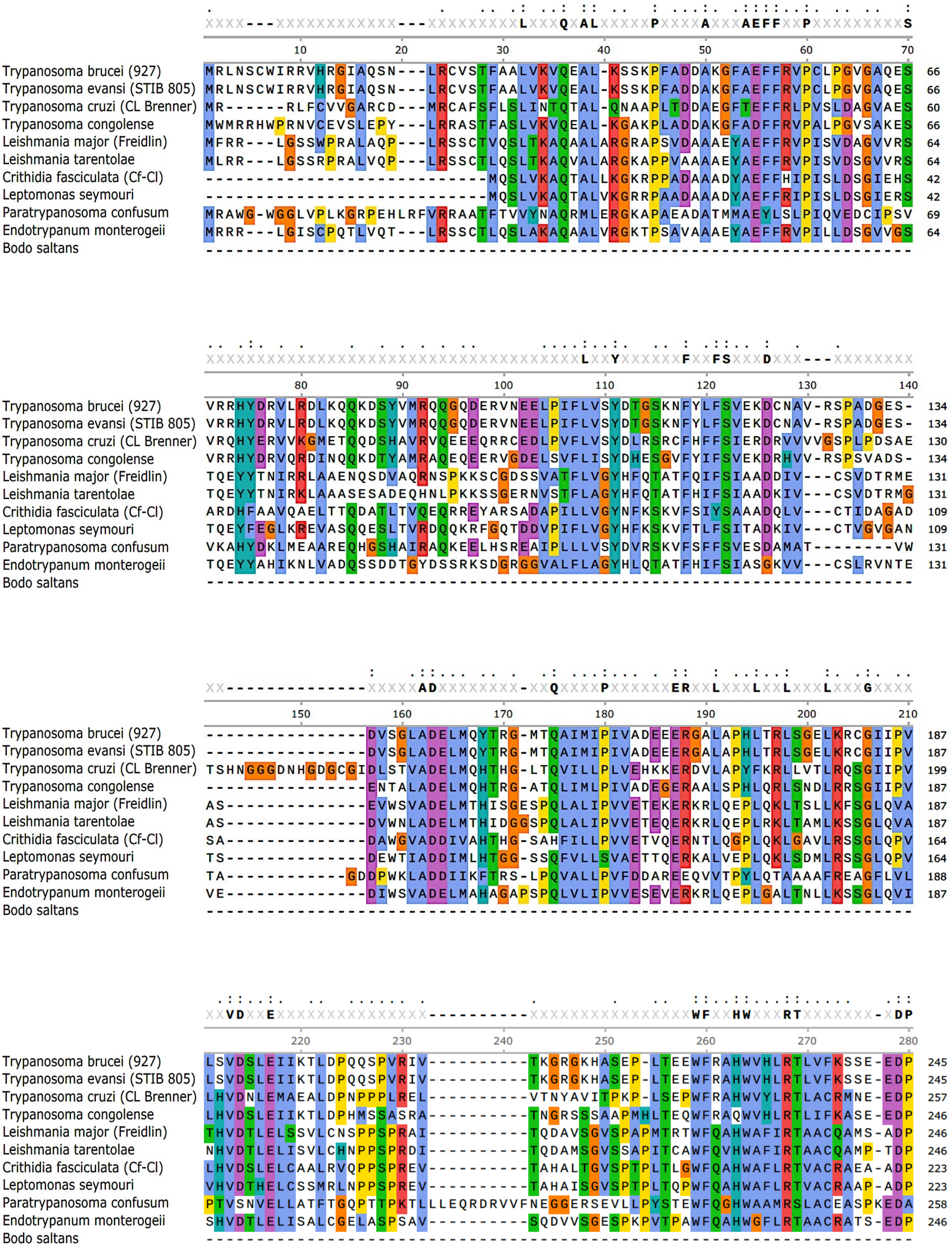
Trypanosoma brucei (927)
Trypanosoma evansi (STIB 805)
Trypanosoma cruzi (CL Brenner)
Trypanosoma congolense
Leishmania major (Freidlin)
Leishmania tarentolae
Crithidia fasciculata (Cf-Cl)
Leptomonas seymouri
Paratrypanosoma confusum
Endotrypanum monterogeii
Bodo saltans

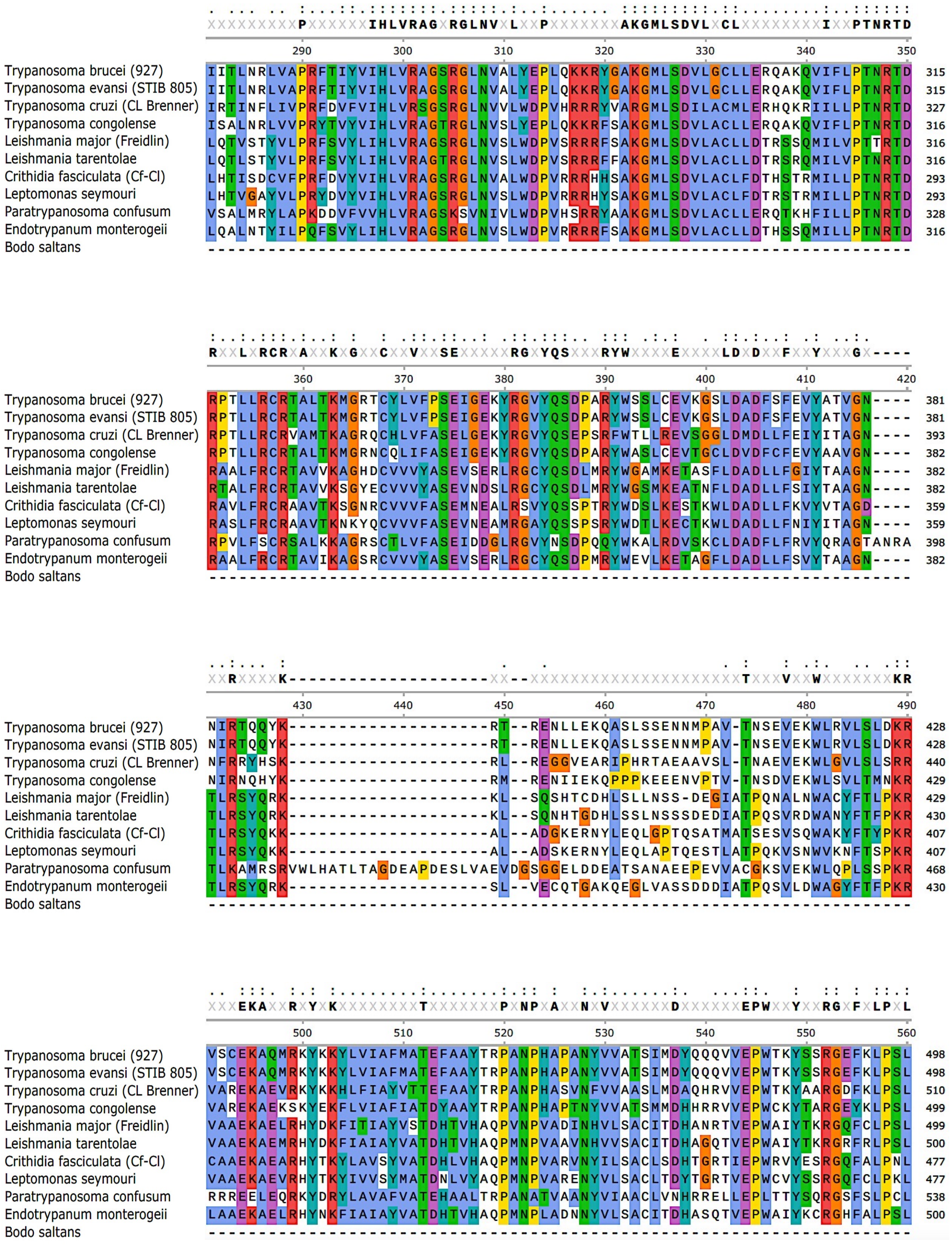
Trypanosoma brucei (927)
Trypanosoma evansi (STIB 805)
Trypanosoma cruzi (CL Brenner)
Trypanosoma congolense
Leishmania major (Freidlin)
Leishmania tarentolae
Crithidia fasciculata (Cf-Cl)
Leptomonas seymouri
Paratrypanosoma confusum
Endotrypanum monterogeii
Bodo saltans
290
300
310
320
330
340
350
360
370
380
390
400
410
420
430
440
450
460
470
480
490
500
510
520
530
540
550
560

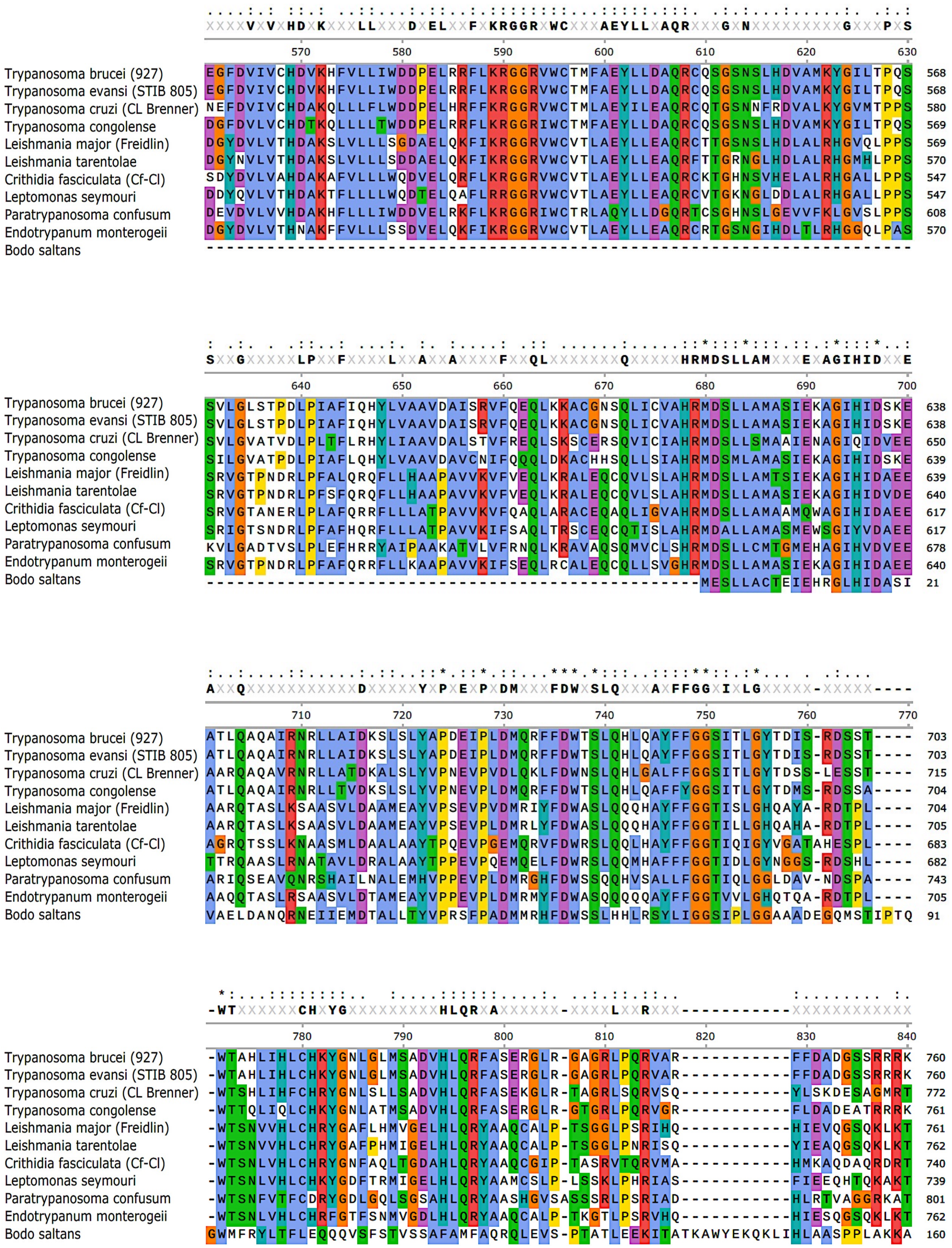
XXXXVXVXHDXKXXXLLXXXDXELXXFXKRGGRXWCXXXAEYLLXAQRXXXGXNXXXXXXXXXGXXXPXS
Trypanosoma brucei (927) EGFDVIVCHDVKHFVLLIWDDPELRRFLKRGGRVWCTMFAEYLLDAQRCQSGSNSLHDVAMKYGILTPQS 568
Trypanosoma evansi (STIB 805) EGFDVIVCHDVKHFVLLIWDDPELRRFLKRGGRVWCTMFAEYLLDAQRCQSGSNSLHDVAMKYGILTPQS 568
Trypanosoma cruzi (CL Brenner) NEFDVIVCHDAKQLLLLFWDDPELHRFFKRGGRIWCTMLAEYILEAQRCQTGSNNFRDVALKYGVMTPPS 580
Trypanosoma congolense DGFDVLVCHDTKQLLLLTWDDPELRRFLKRGGRIWCTMFAEYLLDAQRCQSGSNSLHDVAMKYGILTPQS 569
Leishmania major (Freidlin) DGYDVLVTHDAKSLVLLLSGDAELQKFIKRGGRVWCVTLAEYLLEAQRCTTGSNSLHDLALRHGVQLPPS 569
Leishmania tarentolae DGYNVLVTHDAKSLVLLLSDDAELQKFIKRGGRVWCVTLAEYLLEAQRFTTGRNGLHDLALRHGMHLPPS 570
Crithidia fasciculata (Cf-Cl) SDYDVLVAHDAKAFVLLLWQDVELQRFLKRGGRVWCVTLAEYLLEAQRCKTGHNSVHELALRHGALLPPS 547
Leptomonas seymouri DDYQVLVTHDAKTFLLLLWQDTELQAFLRRGGRVWCVTLAEYLLEAQRCVTGKNGLDDLALRHGALLPPS 547
Paratrypanosoma confusum DEVDVLVVHDAKHFLLLIWDDVELRKFLKRGGRIWCTRLAQYLLDGQRTCSGHNSLGEVVFKLGVSLPPS 608
Endotrypanum monterogeii DGYDVLVTHNAKFFVLLLSSDVELQKFIKRGGRVWCVTLAEYLLEAQRCRTGSNGIHDLTLRHGGQLPAS 570
Bodo saltans ----------------------------------------------------------------------
SXXGXXXXXLPXXFXXXXLXXAXXAXXXXFXXQLXXXXXXXQXXXXXHRMDSLLAMXXXEXAGIHIDXXE
Trypanosoma brucei (927) SVLGLSTPDLPIAFIQHYLVAAVDAISRVFQEQLKKACGNSQLICVAHRMDSLLAMASIEKAGIHIDSKE 638
Trypanosoma evansi (STIB 805) SVLGLSTPDLPIAFIQHYLVAAVDAISRVFQEQLKKACGNSQLICVAHRMDSLLAMASIEKAGIHIDSKE 638
Trypanosoma cruzi (CL Brenner) SVLGVATVDLPLTFLRHYLIAAVDALSTVFREQLSKSCERSQVICIAHRMDSLLSMAAIENAGIQIDVEE 650
Trypanosoma congolense SILGVATPDLPIAFLQHYLVAAVDAVCNIFQQQLDKACHHSQLLSIAHRMDSMLAMASIEKAGIHIDSKE 639
Leishmania major (Freidlin) SRVGTPNDRLPFALQRQFLLHAAPAVVKVFVEQLKRALEQCQVLSLAHRMDSLLAMTSIEKAGIHIDAEE 639
Leishmania tarentolae SRVGTPNDRLPFSFQRQFLLHAAPAVVKVFVEQLKRALEQCQVLSLAHRMDSLLAMASIEKAGIHIDVDE 640
Crithidia fasciculata (Cf-Cl) SRVGTANERLPLAFQRRFLLLATPAVVKVFQAQLARACEQAQLIGVAHRMDSLLAMAAMQWAGIHIDAEE 617
Leptomonas seymouri SRIGTSNDRLPFAFHQRFLLLATPAVVKIFSAQLTRSCEQCQTISLAHRMDALLAMASMEWSGIYVDAEE 617
Paratrypanosoma confusum KVLGADTVSLPLEFHRRYAIPAAKATVLVFRNQLKRAVAQSQMVCLSHRMDSLLCMTGMEHAGIHVDVEE 678
Endotrypanum monterogeii SRVGTPNDRLPFAFQRRFLLKAAPAVVKIFSEQLRCALEQCQLLSVGHRMDSLLAMASIEKAGIHIDAEE 640
Bodo saltans -------------------------------------------------MESLLACTEIEHRGLHIDASI 21
AXXQXXXXXXXXXXDXXXXXYXPXEXPXDMXXXFDWXSLQXXXAXFFGGXIXLGXXXXX-XXXXX----
Trypanosoma brucei (927) ATLQAQAIRNRLLAIDKSLSLYAPDEIPLDMQRFFDWTSLQHLQAYFFGGSITLGYTDIS-RDSST---- 703
Trypanosoma evansi (STIB 805) ATLQAQAIRNRLLAIDKSLSLYAPDEIPLDMQRFFDWTSLQHLQAYFFGGSITLGYTDIS-RDSST---- 703
Trypanosoma cruzi (CL Brenner) AARQAQAVRNRLLATDKALSLYVPNEVPVDLQKLFDWNSLQHLGALFFGGSITLGYTDSS-LESST---- 715
Trypanosoma congolense ATLQAQAIRNRLLTVDKSLSLYVPNEVPLDMQRFFDWTSLQHLQAFFYGGSITLGYTDMS-RDSSA---- 704
Leishmania major (Freidlin) AARQTASLKSAASVLDAAMEAYVPSEVPVDMRIYFDWASLQQQHAYFFGGTISLGHQAYA-RDTPL---- 704
Leishmania tarentolae AARQTASLKSAASVLDAAMEAYVPSEVPLDMRLYFDWASLQQQHAYFFGGTILLGHQAHA-RDTPL---- 705
Crithidia fasciculata (Cf-Cl) AGRQTSSLKNAASMLDAALAAYTPQEVPGEMQRVFDWRSLQQLHAYFFGGTIQIGYVGATAHESPL---- 683
Leptomonas seymouri TTRQAASLRNATAVLDRALAAYTPPEVPQEMQELFDWRSLQQMHAFFFGGTIDLGYNGGS-RDSHL---- 682
Paratrypanosoma confusum ARIQSEAVQNRSHAILNALEMHVPPEVPLDMRGHFDWSSQQHVSALLFGGTIQLGGLDAV-NDSPA---- 743
Endotrypanum monterogeii AAQQTASLRSAASVLDTAMEAYVPPEVPLDMRMYFDWASQQQQQAYFFGGTVVLGHQTQA-RDTPL---- 705
Bodo saltans VAELDANQRNEIIEMDTALLTYVPRSFPADMMRHFDWSSLHHLRSYLIGGSIPLGGAAADEGQMSTIPTQ 91
-WTXXXXXXCHXYGXXXXXXXXXHLQRXAXXXXXX-XXXXLXXRXXX-----------XXXXXXXXXXXX
Trypanosoma brucei (927) -WTAHLIHLCHKYGNLGLMSADVHLQRFASERGLR-GAGRLPQRVAR-----------FFDADGSSRRRK 760
Trypanosoma evansi (STIB 805) -WTAHLIHLCHKYGNLGLMSADVHLQRFASERGLR-GAGRLPQRVAR-----------FFDADGSSRRRK 760
Trypanosoma cruzi (CL Brenner) -WTSHLIHFCHRYGNLSLLSADVHLQRFASEKGLR-TAGRLSQRVSQ-----------YLSKDESAGMRT 772
Trypanosoma congolense -WTTQLIQLCHKYGNLATMSADVHLQRFASERGLR-GTGRLPQRVGR-----------FLDADEATRRRK 761
Leishmania major (Freidlin) -WTSNVVHLCHRYGAFLHMVGELHLQRYAAQCALP-TSGGLPSRIHQ-----------HIEVQGSQKLKT 761
Leishmania tarentolae -WTSNVVHLCHRYGAFPHMIGELHLQRYAAQCALP-TSGGLPNRISQ-----------YIEAQGSQKLKT 762
Crithidia fasciculata (Cf-Cl) -WTSNLVHLCHRYGNFAQLTGDAHLQRYAAQCGIP-TASRVTQRVMA-----------HMKAQDAQRDRT 740
Leptomonas seymouri -WTSNLVHLCHKYGDFTRMIGELHLQRYAAMCSLP-LSSKLPHRIAS-----------FIEEQHTQKAKT 739
Paratrypanosoma confusum -WTSNFVTFCDRYGDLGQLSGSAHLQRYAASHGVSASSSRLPSRIAD-----------HLRTVAGGRKAT 801
Endotrypanum monterogeii -WTSNLVHLCHRFGTFSNMVGDLHLQRYAAQCALP-TKGTLPSRVHQ-----------HIESQGSQKLKT 762
Bodo saltans GWMFRYLTFLEQQQVSFSTVSSAFAMFAQRQLEVS-PTATLEEKITATKAWYEKQKLIHLAASPPLAKKA 160

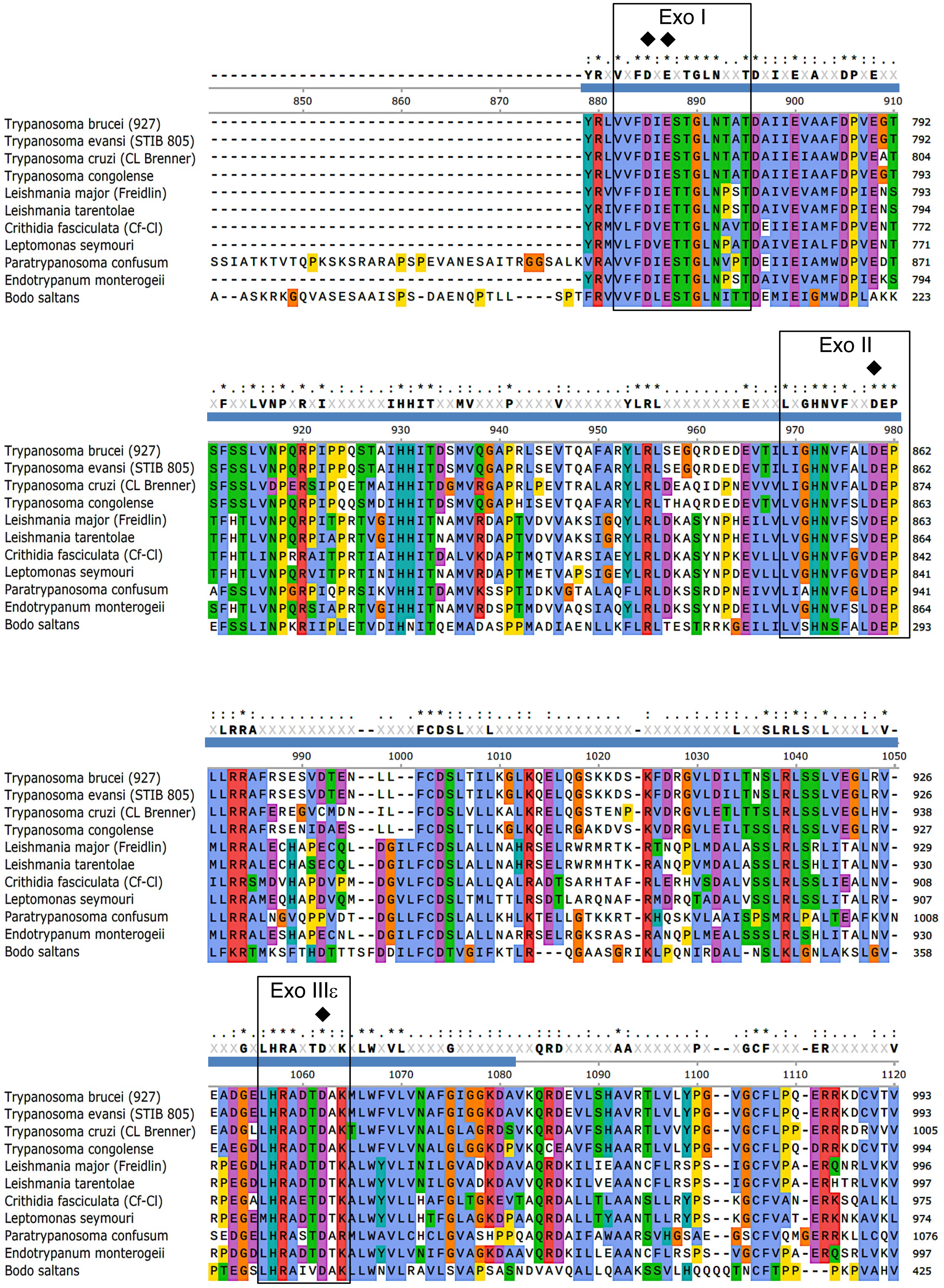

Exo I
Exo II
Exo IIIε
Trypanosoma brucei (927)
Trypanosoma evansi (STIB 805)
Trypanosoma cruzi (CL Brenner)
Trypanosoma congolense
Leishmania major (Freidlin)
Leishmania tarentolae
Crithidia fasciculata (Cf-Cl)
Leptomonas seymouri
Paratrypanosoma confusum
Endotrypanum monterogeii
Bodo saltans

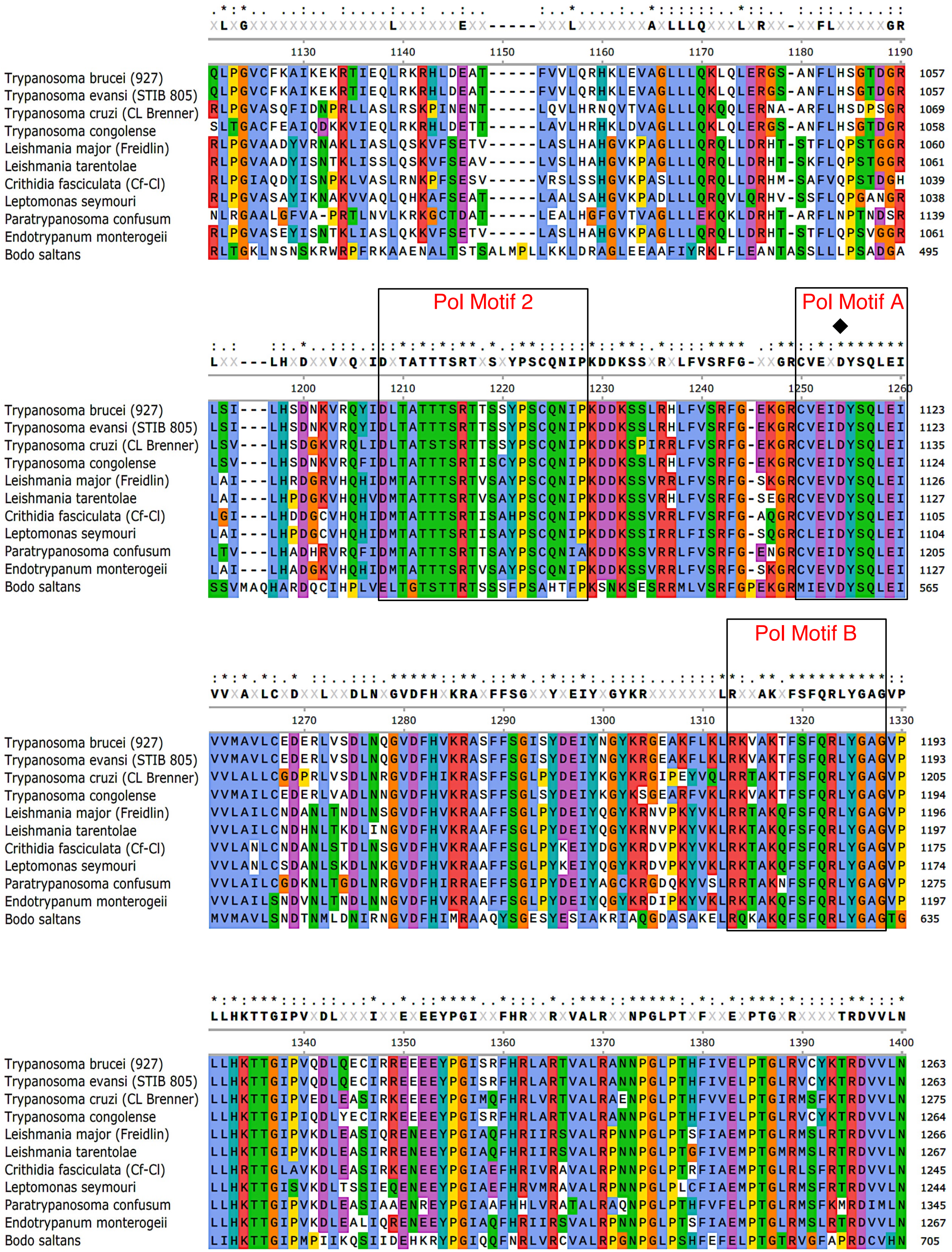
Pol Motif 2
Pol Motif A
Pol Motif B
Trypanosoma brucei (927)
Trypanosoma evansi (STIB 805)
Trypanosoma cruzi (CL Brenner)
Trypanosoma congolense
Leishmania major (Freidlin)
Leishmania tarentolae
Crithidia fasciculata (Cf-Cl)
Leptomonas seymouri
Paratrypanosoma confusum
Endotrypanum monterogeii
Bodo saltans

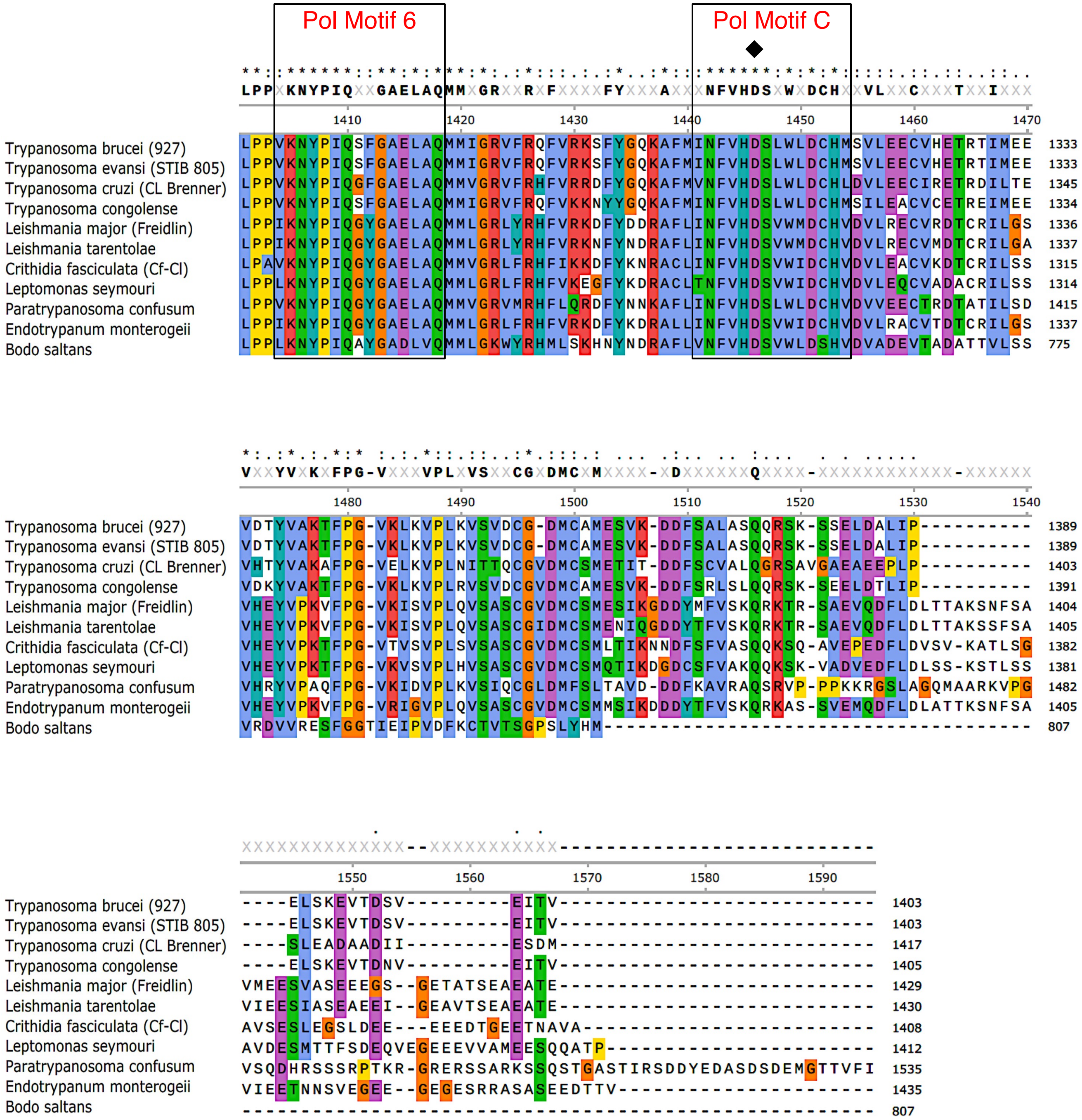


**Figure S5. Alignment of Kinetoplastid POLIB sequences**. The aligned POLIB sequences are: *Trypanosoma brucei brucei, Trypanosoma evansi*, *Trypanosoma cruzi*, *Trypanosoma congolense*, *Leishmania major, Leishmania tarentolae*, *Crithidia fasciculata*, *Leptomonas seymouri*, *Paratrypanosoma confusum*, *Endotrypanum monterogeli* and *Bodo saltans*. Coloring is based on Clustal X formatting. Concensus sequence was generated with a threshold >75%. Highly conserved and invariant residues are small circles and asterisks, respectively. Diamonds indicate the catalytic aspartic or glutamic acid residue. Black squares indicate five of the six most conserved regions in family A DNA polymerases. Blue bar indicates the thumb insertion with sequence similarity to DEDDh 3'-5' exonucleases.

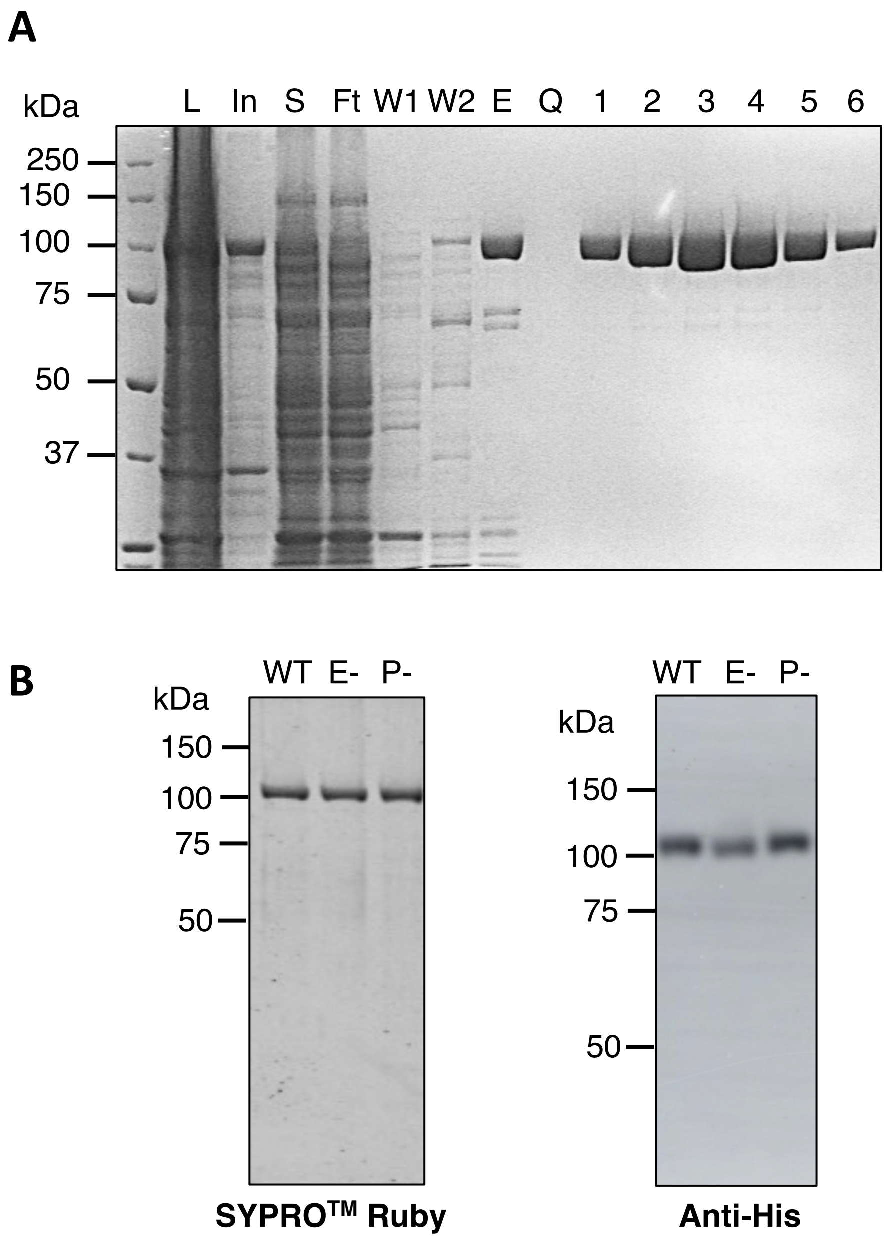


**Figure S6. Preparation of recombinant TbPOLIB variants. (A)** Coomasie stained 8% SDS-PAGE gel of IBWT purification progression. L; cell lysate, In; insoluble fraction, S; soluble fraction, Ft; flowthrough of the Ni column, W1; wash with lysis buffer, W2; wash with wash buffer, E; elutate from Nickel column, Q; flowthrough from the Q column, 1-6; selected fractions from Q column NaCl gradient elution. **(B)** 500 ng each POLIB variant was analyzed on 8% SDS-PAGE gel and either stained with Sypro ruby (left) or transfered to membrane and detected with anti-His antibody (right). WT, wild type variant; E-, exonuclease deficient variant; P-, polymerase deficient variant.

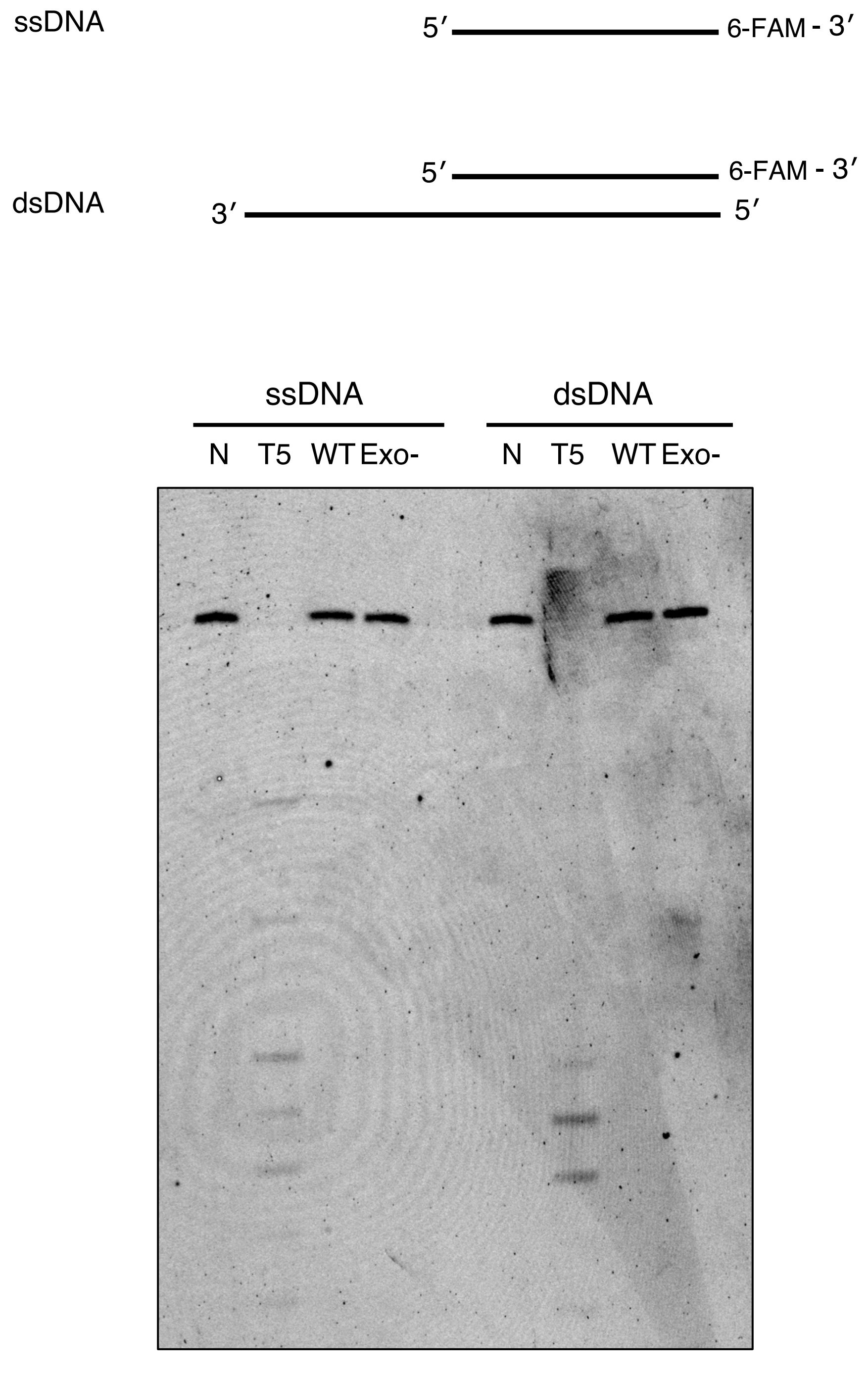


**Figure S7. POLIB Lacks 5′-3′ Activity on DNA substrates.** 200 nM POLIB variant was used to initiate each reaction using standard conditions (pH 7.0, 5 mM $MgCl_2$). T5 exonuclease (10U), positive control. All reactions were quenched after 30 minutes. Negative controls contain no enzyme and were quenched at time 0. dsDNA, 3′ FAM labeled 22mer annealed to the unlabeled 42mer template; ssDNA, 3′ FAM labeled 22mer. WT, wild type variant; Exo-, exonuclease deficient variant.

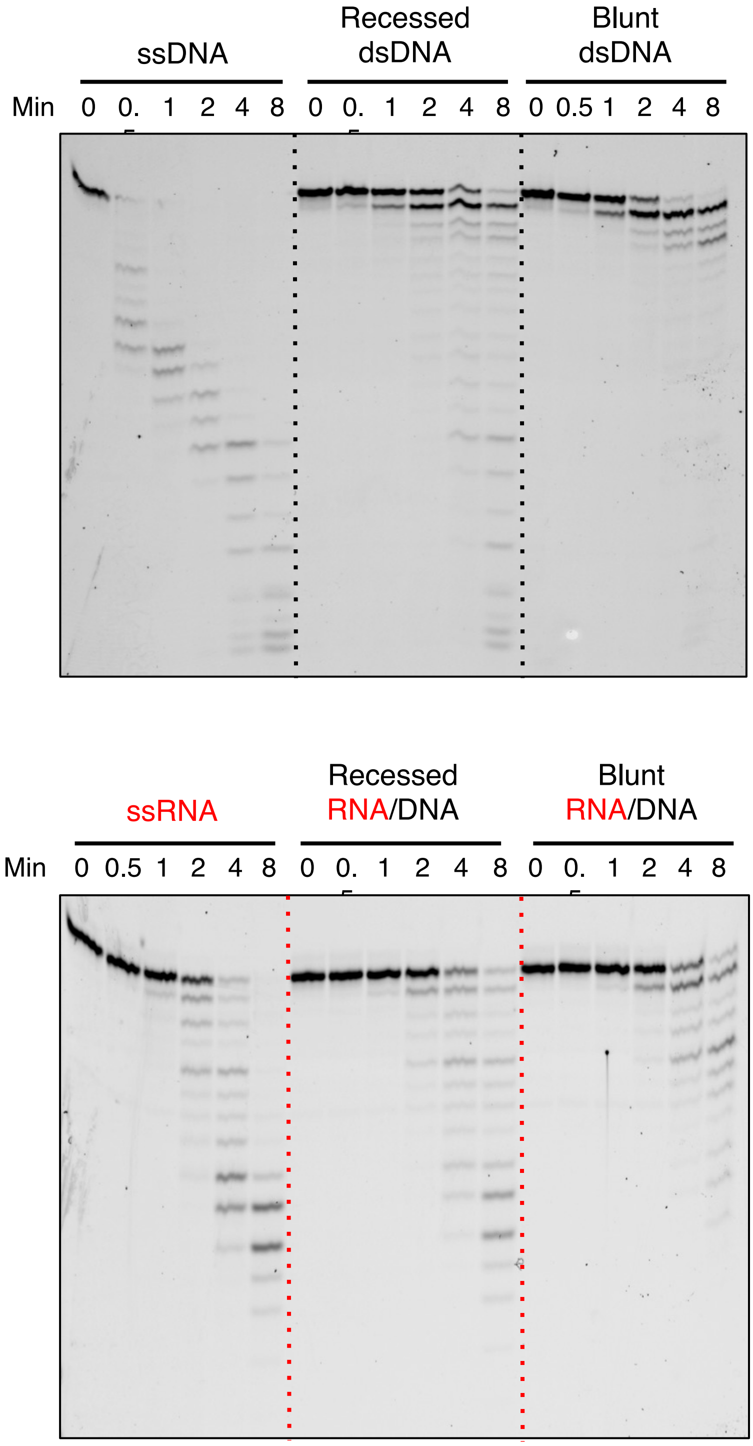


**Figure S8. Representative images used for quantification of data in Figure 3.**

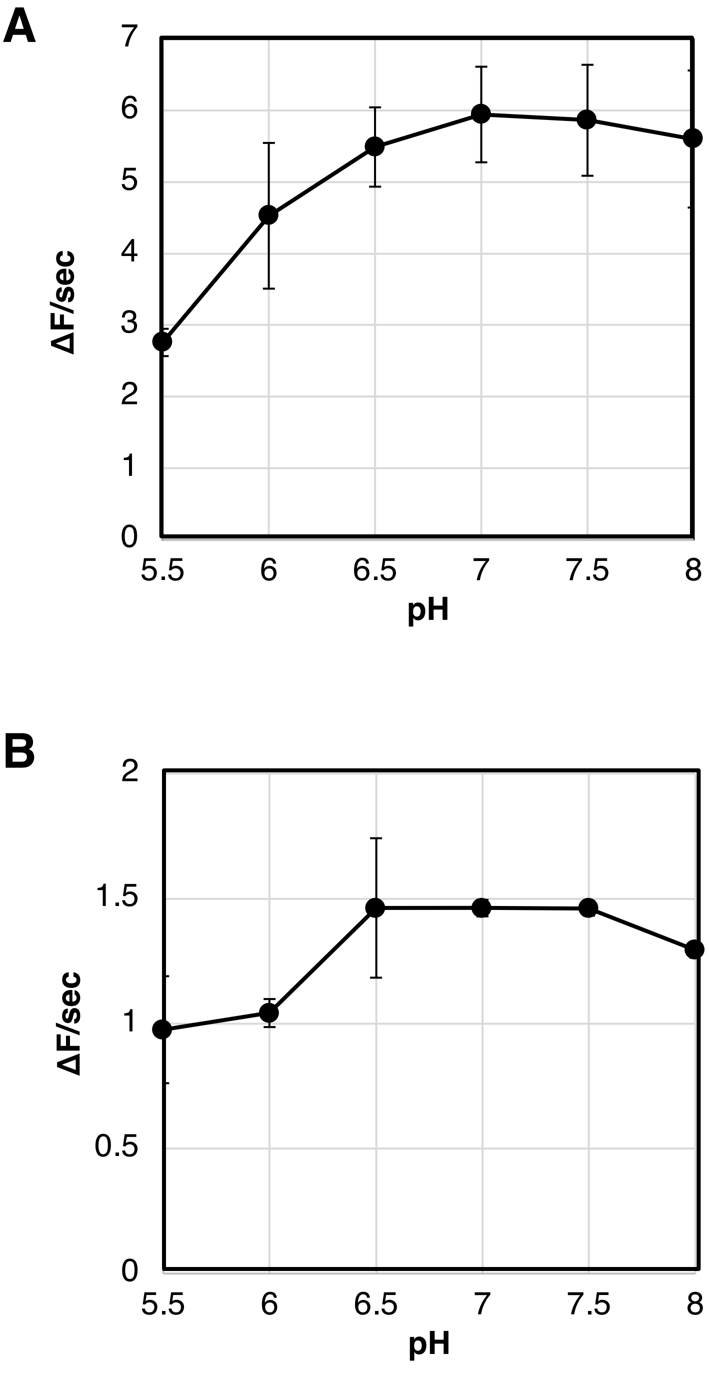


**Figure S9. Rate of POLIB exonuclease activity on A) ssDNA and B) blunt dsDNA.** Each 2AP excision assay contained standard buffer conditions, 200 nM IBWT and 5 μM 2AP position 1. Only the pH varied as indicated. Error bars represent standard deviation of three replicates, some error bars are too small to be visible on this chart.

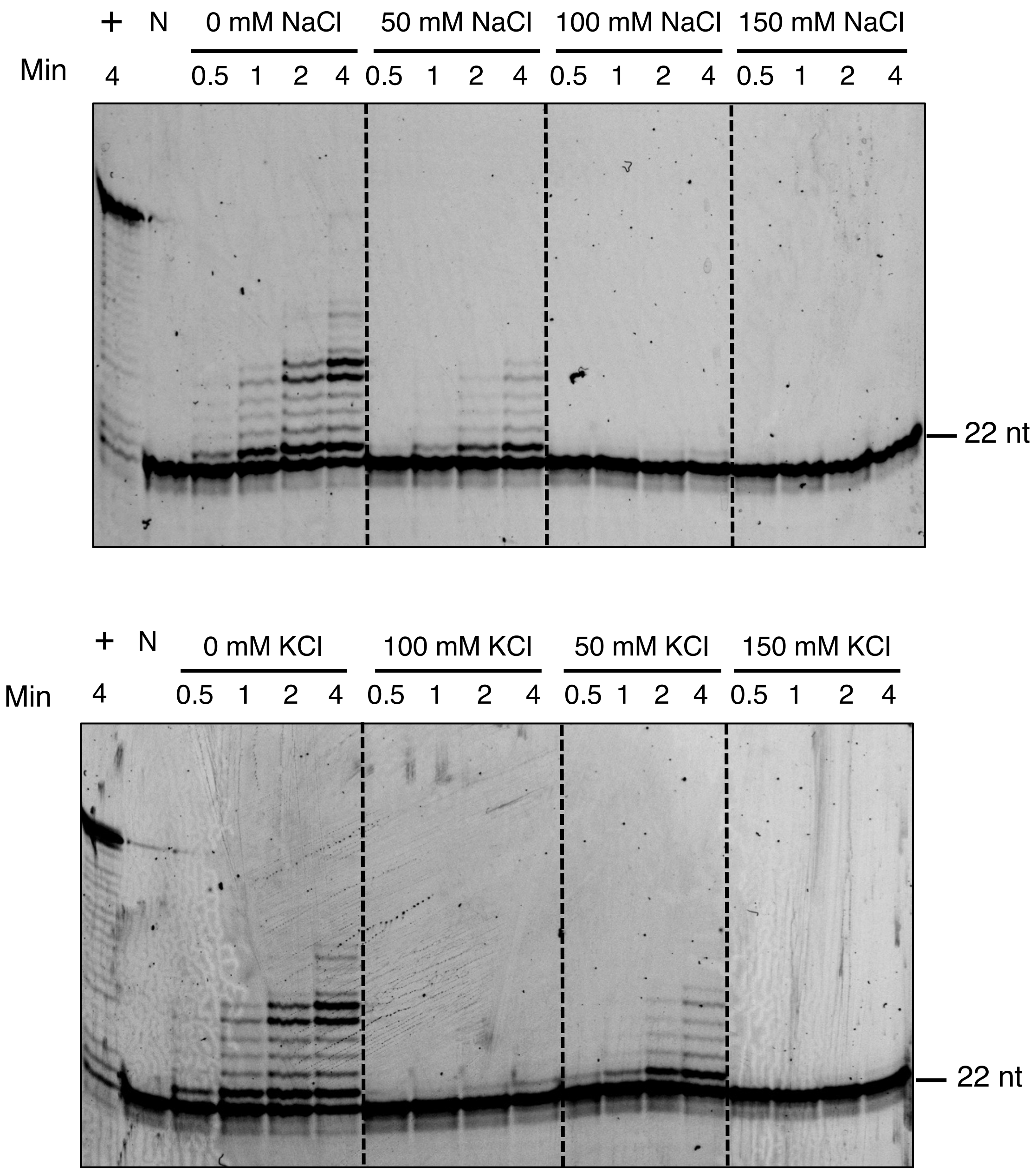


**Figure S10. Salt optimum of primer extension activity for IBexo-.** Reactions contained standard rection buffer (pH 7.0), varying salt concentrations as indicated and was initiated with 200 nM IBExo-. Positive control (+), 1U Klenow pH 8.0, no salt in buffer. Negative control (N) quenched at time 0.

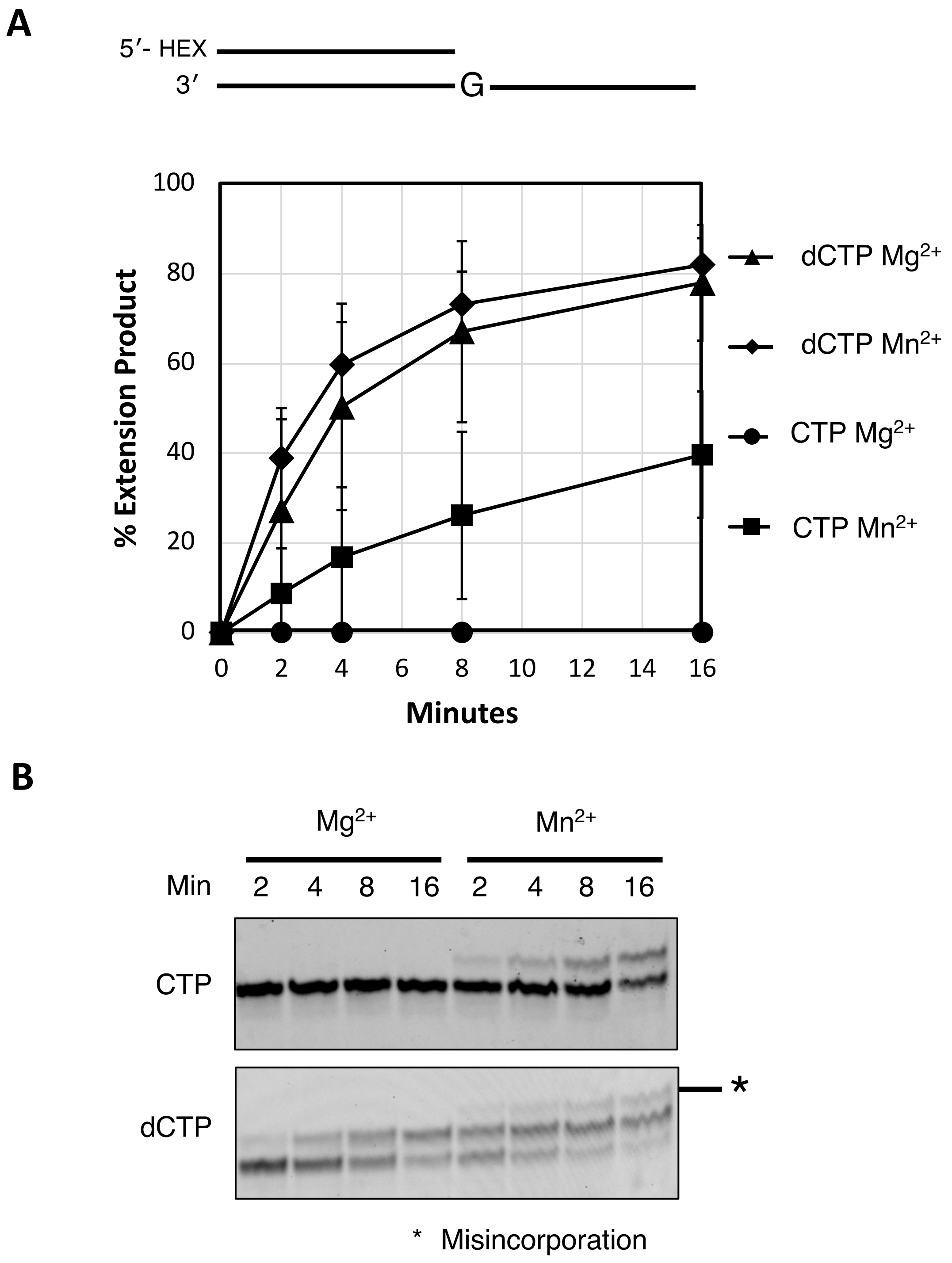


**Figure S11. POLIB incorporates nucleotides more rapidly with $Mn^{2+}$ than $Mg^{2+}$.** Reactions include 200 nM IBExo- 5 mM CTP or dCTP, 50 mM Tris pH 7.0, 1 mM DTT, 0.1% BSA, 8.4 µM DNA, reactions started with 500 M $MgCl_2$ or 5 mM $MnCl_2$ A) quantification of % primer extended across three replicates with different POLIB preparations. B) Representative of primer extension reactions.

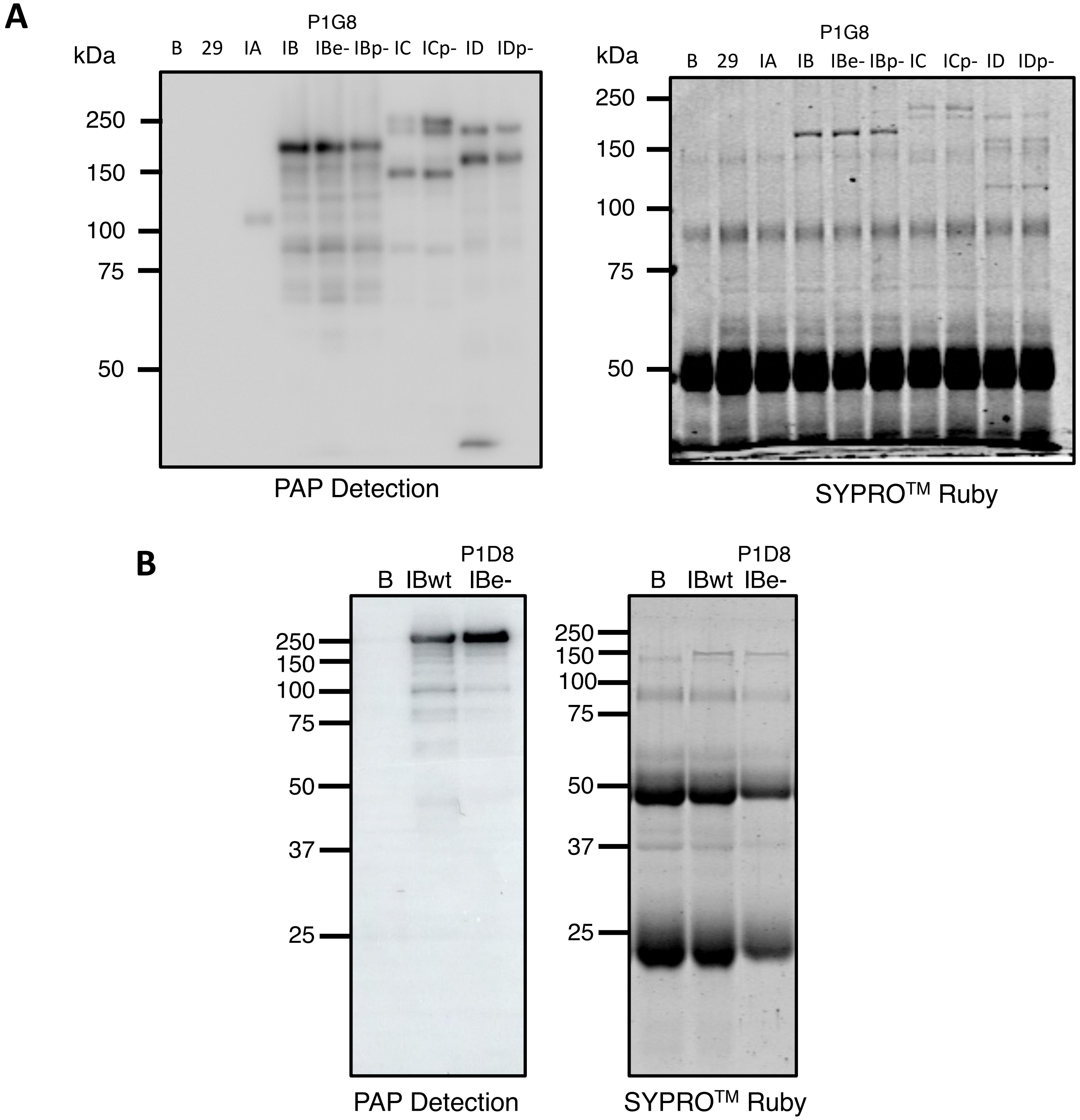


**Figure S12. SDS-PAGE and Western Blots of PTP-tagged Pol I-like paralog variants immunoprecipitated from *T. brucei* procyclic cells.** **(A)** Detection of Pol I-like variants on IgG Sepharose beads used in primer extension assay Figure 7A. PAP detection or SYPRO$^{TM}$ Ruby for total protein. B; Beads-only, 29; IP with 29-13 procyclic cells, IA; IA$^{MHTAP}$, IB; IBwt$^{PTP}$, IBe-; IBexo-$^{PTP}$ (clone P1G8), IBp-; IBpol-$^{PTP}$, IC; ICwt$^{PTP}$, ICp-; ICpol-$^{PTP}$, ID; IDwt$^{PTP}$, IDp-; IDpol$^{PTP}$. **(B)** Representative PAP and SYPRO$^{TM}$ Ruby detection for IP samples of IBwt$^{PTP}$ and IBexo-$^{PTP}$ (clone P1D8) used in Figure 7B. Labels as in A.

### A. POLIB Codon-Optimized Nucleotide Sequence

AGCGAAGTTGAAAAATGGCTGCGTGTTCTGAGCCTGGATAAACGTGTTAGCTGTGAAAAAGCACAGATGCGCAAATACAAA
AAGTATCTGGTGATTGCATTTATGGCCACCGAATTTGCAGCATATACCCGTCCGGCAAATCCGCATGCACCGGCAAATTAT
GTTGTTGCAACCAGCATTATGGATTACCAGCAGCAGGTTGTTGAACCGTGGACCAAATATAGCAGCCGTGGTGAATTTAAA
CTGCCGAGCCTGGAAGGTTTTGATGTTATTGTTTGCCATGACGTGAAACACTTTGTTCTGCTGATTTGGGATGATCCGGAA
CTGCGTCGTTTTCTGAAACGTGGTGGTCGTGTTTGGTGTACCATGTTTGCAGAATATCTGCTGGATGCACAGCGTTGTCAG
AGCGGTAGCAATAGCCTGCATGATGTTGCAATGAAATATGGTATTCTGACACCGCAGAGCAGCGTTCTGGGTCTGAGCACA
CCGGATCTGCCGATTGCCTTTATTCAGCATTATCTGGTTGCAGCAGTTGATGCAATTAGCCGTGTTTTTCAAGAGCAGCTG
AAAAAAGCATGTGGTAATAGCCAGCTGATTTGTGTTGCACATCGTATGGATAGCCTGCTGGCAATGGCAAGCATTGAAAAA
GCCGGTATTCATATCGATAGCAAAGAAGCAACCCTGCAGGCACAGGCAATTCGTAATCGTCTGCTGGCCATTGATAAATCA
CTGAGCCTGTATGCACCGGATGAAATTCCGCTGGATATGCAGCGTTTTTTTGATTGGACCAGCCTGCAGCATCTGCAAGCA
TATTTCTTTGGTGGTAGCATTACCCTGGGTTATACCGATATTAGTCGTGATAGCAGCACCTGGACCGCACATCTGATTCAT
CTGTGTCATAAATATGGCAATCTGGGCCTGATGAGCGCAGATGTTCATCTGCAGCGCTTTGCAAGCGAACGTGGTCTGCGT
GGTGCAGGTCGTCTGCCGCAGCGTGTTGCACGCTTTTTTGATGCAGATGGTAGCAGTCGTCGTAAATATCGTCTGGTT
GTGTTTGATATTGAAAGCACCGGTCTGAATACCGCAACCGATGCCATTATTGAAGTTGCAGCCTTTGATCCGGTTGAAGGC
ACCAGCTTTAGCAGCCTGGTTAATCCGCAGCGTCCGATTCCGCCTCAGAGCACCGCAATTCATCATATTACCGATAGCATG
GTTCAGGGTGCACCGCGTCTGAGCGAAGTTACCCAGGCATTTGCACGTTATCTGCGCCTGAGTGAAGGTCAGCGTGATGAA
GATGAAGTTACCATTCTGATTGGCCATAATGTTTTTGCACTGGATGAACCGCTGCTGCGTCGTGCATTTCGTAGCGAAAGC
GTTGATACCGAAAATCTGCTGTTTTGTGATAGCCTGACAATTCTGAAAGGCCTGAAACAAGAACTGCAGGGTAGCAAAAAA
GATAGCAAATTTGATCGTGGCGTGCTGGATATTCTGACCAATAGTCTGCGTCTGAGTAGCCTGGTTGAAGGTCTGCGCGTT
GAAGCCGATGGTGAACTGCATCGTGCAGATACCGATGCAAAAATGCTGTGGTTTGTTCTGGTTAATGCCTTTGGTATTGGT
GGTAAAGATGCAGTTAAACAGCGTGACGAAGTGCTGAGCCATGCAGTTCGTACCCTGGTTCTGTATCCTGGTGTTGGTTGT
TTTCTGCCGCAAGAACGTCGTAAAGATTGTGTTACCGTTCAGCTGCCTGGTGTTTGTTTTAAAGCCATTAAAGAAAAACGC
ACCATTGAGCAGCTGCGTAAACGTCATCTGGATGAAGCAACCTTTGTTGTTCTGCAGCGTCATAAACTGGAAGTTGCCGGT
CTGCTGCTGCAGAAACTGCAGCTGGAACGTGGTAGCGCAAATTTTCTGCATAGCGGCACCGATGGTCGTCTGAGCATTCTG
CATTCAGATAATAAAGTGCGCCAGTATATTGATCTGACCGCAACCACCACCAGTCGTACCACCAGTAGCTATCCGAGCTGC
CAGAATATTCCGAAAGATGATAAAAGCAGCCTGCGTCACCTGTTTGTTAGCCGTTTTGGTGAAAAAGGTCGTTGCGTGGAA
ATTGATTATTCCCAGCTGGAAATTGTTGTTATGGCAGTTCTGTGCGAAGATGAACGTCTGGTTAGCGATCTGAATCAGGGT
GTTGATTTTCATGTTAAACGTGCCAGCTTTTTTAGCGGCATTAGCTATGATGAAATCTATAACGGTTATAAACGCGGTGAA
GCCAAATTCCTGAAACTGCGTAAAGTTGCAAAAACCTTTAGCTTTCAGCGTCTGTATGGTGCCGGTGTTCCTCTGCTGCAT
AAAACCACCGGTATTCCGGTTCAGGATCTGCAAGAATGTATTCGTCGTGAAGAAGAATATCCGGGTATTTCACGTTTT
CATCGTCTGGCACGTACCGTTGCACTGCGTGCAAATAATCCGGGTCTGCCGACACATTTTATTGTTGAACTGCCGACAGGC
CTGCGTGTTTGCTATAAAACCCGTGATGTTGTGCTGAATCTGCCTCCGGTTAAAAACTATCCGATTCAGAGCTTTGGTGCA
GAACTGGCACAGATGATGATTGGTCGTGTGTTTCGTCAGTTTGTGCGTAAAAGTTTCTATGGCCAGAAAGCCTTTATGATC
AACTTTGTGCATGATAGTCTGTGGCTGGATTGTCATATGAGTGTTCTGGAAGAATGTGTTCATGAAACCCGTACCATTATG
GAAGAGGTTGATACCTATGTGGCCAAAACCTTTCCGGGTGTTAAACTGAAAGTGCCGCTGAAAGTTAGCGTTGATTGTGGT
GTTGATATGTGTGCAATGGAAAGCGTGAAAGATGACTTTAGCGCACTGGCAAGCCAGCAGCGTAGCAAATCAAGCGAACTG
GATGCCCTGATTCCTGAACTGTCAAAAGAAGTGACCGATAGCGTTTAA

### B. Oligonucleotides used for Site Directed Mutagenesis

| Mutagenic site | Sequence (5'-3') |
|---|---|
| Pol Domain D1117A | ATTTCCAGCTGGGAATAA**G**CAATTTCCACGCAACGAC |
| Pol Domain D1309A | CAATCCAGCCACAGACTA**G**CATGCACAAAGTTGATCA |
| Exo Domain D767N, E769Q | CAGACCGGTGCTTT**G**AATAT**T**AAACACAACCAGACGATATTTACGA |

**Table S1. Nucleotide reagents used for POLIB recombinant constructs. A.** recoded region corresponding to POLIB nucleotide optimized for E. coli expression. **B**. Primers for site-directed mutagenesis. Mutagenic sites are in **red bold**.

#### A. Epitope tagged trypanosome cell lines for this study

| Cell Line | Mutation |
|---|---|
| POLIA$^{MHTAP}$ (IAwt) | none |
| POLIB$^{PTP}$ (IBwt) | none |
| POLIBExo-$^{PTP}$ (IBe-) | D767N, E769Q |
| POLIBPol-$^{PTP}$ (IBp-) | D1117A, D1309A |
| POLIC$^{PTP}$ (ICwt) | none |
| POLICPol-$^{PTP}$ (ICp-) | D1380A, D1592A |
| POLID$^{PTP}$ (IDwt) | none |
| POLIDPol-$^{PTP}$ (IDp-) | D1346A, D1565A |

#### B. Oligonucleotides used for generating trypanosomes cell lines

| Name | Sequence (5'-3') | Purpose |
|---|---|---|
| POLIA F | TTATTAGGATCCATGTCGTTACATGGTCATC | Subcloning |
| POLIA R | TTATTAGGATCCACTCGGCACCCGCAGTC | |
| POLIB F | GTAACTCGAGATGCGGCTAAATAGCTGC | |
| POLIB R | ACACTCTAGACACCGTAATTTCTACACTGTCAG | |
| POLID F | TTATTACAATTGATGCTGCGGCGGCTCTTC | |
| POLID R | TTATTAGGATCCCAGTGTCTCCTCAATGACAACGG | |
| POLIB Rc F | gcttcaattgctcgaGAGATGCGGCTAAATAGCTGC | Gibson |
| POLIB Rc R | gtaccgggccggatcCGACTGATATACCCCACGATAC | |
| POLIB M1 | GGCCGTTGCGTGGAAATTGC**C**TACTCGCAGCTG | Mutagenesis |
| POLIB M2 | GATAAACTTCGTACACG**C**TTCGCTTTGGCTCGAC | |
| POLIB M3 | GCGCAAGTACAGACTAGTTGTATTT**A**ATATT**C**AATCCACGGGATTG | |
| POLID M1 | GGGAATGTGTATTGAGGCAG**C**TTATTCACAGCTTGAAGTCG | |
| POLID M2 | ACTAATCAACACCGTGCACG**C**CTGCGTATGGATTGACTG | |
| POLID M3 | ACTCAAAGGCGGTAATACGGTTATCCACAGAATCAGG | |
| POLID M4 | CCAGTGAATTGTAATACGACTCACTATAGGGCGAATTGG | |

**Table S2. Generation of epitope tagged *T. brucei* Pol I-like paralogs for overexpression and immunoprecipitation.** Underline, restriction enzyme sites, small case, vector backbone sequence; **Red bold**, mutagenic sites.
